# Association Mapping Across a Multitude of Traits Collected in Diverse Environments Identifies Pleiotropic Loci in Maize

**DOI:** 10.1101/2022.02.25.480753

**Authors:** Ravi V. Mural, Guangchao Sun, Marcin Grzybowski, Michael C. Tross, Hongyu Jin, Christine Smith, Linsey Newton, Carson M. Andorf, Margaret R. Woodhouse, Addie M. Thompson, Brandi Sigmon, James C. Schnable

## Abstract

Classical genetic studies have identified many cases of pleiotropy where mutations in individual genes alter many different phenotypes. Quantitative genetic studies of natural genetic variants frequently examine one or a few traits, limiting their potential to identify pleiotropic effects of natural genetic variants. Widely adopted community association panels have been employed by plant genetics communities to study the genetic basis of naturally occurring phenotypic variation in a wide range of traits. High-density genetic marker data – 18M markers – from two partially overlapping maize association panels comprising 1,014 unique genotypes grown in field trials across at least seven US states and scored for 162 distinct trait datasets enabled the identification of of 2,154 suggestive marker-trait associations and 697 confident associations in the maize genome using a resampling-based genome-wide association strategy. The precision of individual marker-trait associations was estimated to be three genes based a reference set of genes with known phenotypes. Examples were observed of both genetic loci associated with variation in diverse traits (e.g. above-ground and below-ground traits), as well as individual loci associated with the same or similar traits across diverse environments. Many significant signals are located near genes whose functions were previously entirely unknown or estimated purely via functional data on homologs. This study demonstrates the potential of mining community association panel data using new higher density genetic marker sets combined with resampling-based genome-wide association tests to develop testable hypotheses about gene functions, identify potential pleiotropic effects of natural genetic variants, and study genotype by environment interaction.

## Introduction

Association mapping, initially on a gene-by-gene level and later at a genome-wide scale, has been widely adopted as a tool to identify natural genetic variants controlling variation in both quantitative and qualitative traits. In the plant genetics community, logistical and scientific constraints have driven the development and widespread adoption of community association panels comprising sets of distinct plant genotypes which can be propagated and shared, whether through the use of homozygous inbred lines or clonal propagation. In the earlier era where association mapping was conducted on a gene-by-gene level, the use of community association panels allowed the work of estimating population structure within the population to be conducted once rather than for each independent study. In the later era of genome-wide association studies, the use of community association panels again provided substantial practical benefits: the time-consuming and expensive process of genotyping hundreds of thousands or millions of genetic markers across hundreds of individuals had to be undertaken only once to enable an effectively infinite number of studies on the genes controlling different traits by different research groups.

The use of community association panels by many independent research groups to investigate diverse research questions results in data on a wide range of individual traits for genetically identical individuals across one or more environments. This provides significant opportunities to investigate both pleiotropy, the effect of a single genetic locus on multiple phenotypes, and genotype by environment interactions, where the same allele influences the same phenotype in different ways in different environments. In addition, associations between a given genetic locus and a particular trait which are only marginally significant in several individual studies can often be assigned a higher degree of confidence when the same association is identified across multiple studies. Finally, marginal statistical signals from genome-wide association studies can assist in the interpretation of later mutant mapping, gene expression, or selection scans, but only if the initial studies results are made available in a method which is easy to capture and cross reference.

In maize, an early widely adopted community association panel was the Maize Association Panel (MAP), also referred to variously as the maize 282 panel and the Buckler-Goodman Association panel (Table S1). MAP initially consisted of 302 diverse inbreds estimated to represent 80% of the genetic diversity within maize, although several of these were dropped in later years based on poor seed increasability or other factors^1,2^. Slow decreases in population size over time are a common feature of many community association panels. Two challenges were observed with the initial maize association panel. Firstly, while the MAP panel captured a large proportion of total maize genetic diversity, it consisted primarily of older public sector lines with limited representation of current temperate elite germplasm. Secondly, many of the MAP lines were difficult to grow and increase in the northern US Corn belt. As a result, two additional panels were generated: 1) The Shoot Apical Meristem association panel (SAM panel) which included many of the MAP genotypes augmented with expired Plant Variety Patent lines generated by the major seed companies in the USA^3^, and 2) The Wisconsin Diversity Panel (WiDiv) developed by selecting non-redundant and diverse genotypes which were able to complete their life cycle and produce significant amounts of seed when grown in Madison, Wisconsin^4^.

As sequencing technologies have improved, new sets of genetic markers have been deployed for existing community association panels with increasing degrees of genetic resolution. The MAP was initially genotyped with a modest number of (<100) SSR markers to estimate population structure as a potential confounder for single gene association tests^2^. The WiDiv panel was initially genotyped with 1,536 microarray-based markers^4^. The SAM panel was initially genotyped using sequencing of mRNA samples from each line, enabling the identification and scoring of 1.2M segregating SNP markers^3^. A new high-density marker set for the WiDiv panel was also generated by sequencing mRNA samples, providing a set of 900k segregating genetic markers in this population^5,6^. The original MAP population which shares many genotypes with both the SAM and WiDiv panels was resequenced as part of the Maize HapMap3 project increasing the number of segregating genetic markers to 83M^7^. A subset of lines from the WiDiv panel were resequenced, resulting in a set of 3.1M SNPs scored across 511 genotypes^8^.

Here we employ a combination of published resequencing data to generate a common set of 18M genetic markers scored across the union of 1014 genotypes present in the SAM and/or WiDiv association panels. We assemble a set of 162 trait datasets which have been scored across different subsets of these 1014 genotypes, including both previously published studies conducted across seven US states and new trait data collected from field trials conducted in Lincoln, Nebraska USA. Using resampling-based genome-wide association studies (GWAS) we define evidence based best practices for mapping intervals around GWAS hits in these maize populations based on either physical distance or gene rank order. We identify suggestive signatures of pleiotropic effects for a number of genetic loci, resulting in a data release of genomic intervals associated with 2,154 confident or suggestive GWAS hits across these 162 trait datasets to aid in the reuse of these trait data in future genomic and genetic studies.

## Materials & Methods

### Collection of Published Trait Data

Papers publishing maize analyses were identified by searching papers citing the initial description of the first iteration of the WiDiv panel^4^, the initial description of the SAM diversity panel^3^, or the expanded WiDiv panel^6^. Screening of studies citing one or more of these papers concluded on 25^th^ June 2021. Published studies were excluded if we were unable to locate de-anonymized trait values for individual accessions or if less than 200 total accessions were phenotyped. If two studies indicated that the same trait was collected from the same lines in the same location in the same year, only one version of the data was retained. If a study published both data from individual environments and aggregated estimates across environments (e.g. averages, Best Linear Unbiased Predictions (BLUPs), or Best Linear Unbiased Estimates (BLUEs)) only individual environment trait data was retained. If only aggregate estimates across environments were published, aggregated traits were employed. After preliminary analysis with MLM-based GWAS several other trait datasets were discarded when it proved impossible to effectively control false discoveries across the genome. The final data file of all accession-level trait values employed in this study is provided as Table S2.

### Trait Data Not Previously Published

A set of 752 maize genotypes, which were a strict subset of the WiDiv panel, and included 254 of 369 genotypes from the SAM diversity panel were evaluated in a field experiment conducted in Lincoln, Nebraska in the summer of 2020. The experimental design of this field experiment has been previously described^9^. Briefly, the field was laid out in a randomized complete block design with two blocks of 840 plots including a repeated check genotype (B97). Each plot was two rows, 7.5 feet (approximately 2.3 meters) long with 30 inch row spacing (approximately 0.76 meters), 4.5 inch spacing between sequential plants (approximately 11.5 centimeters) and 30 inch alleyways between sequential plots (approximately 0.76 meters). The field was planted on May 6^th^, 2020 and was located at the University of Nebraska-Lincoln’s Havelock Farm (40.852 N, 96.616 W).

Tassel architecture phenotypes were collected once tassels had fully emerged for three randomly selected plants per plot, avoiding edge plants. Tassel lengths were measured from the basal primary tassel branch to the tip of the tassel spike. Branch zone length was defined as the length from the basal primary tassel branch to the top primary tassel branch. Tassel spike length was defined as the length from the top primary tassel branch to the tip of the tassel spike. The total number of primary tassel branches were also counted as well as the number of these primary tassel branches that were initiated, but later aborted (Figure S1).

Male and female flowering times for each plot were scored on the first day that 50% of plants has visible pollen shed or visible silks, respectively. Root and stalk lodging were scored at the end of the growing season as a percent of extant plants in the plot, following the published Genomes to Fields phenotyping protocol for both traits^10^. Leaf phenotypes – leaf length, leaf width, and leaf angle – were measured for two plants per plot and collected from each plot after anthesis and silking. One plant was randomly selected from each of the two rows for measurement, avoiding edge plants when possible. Leaf length was measured from the leaf ligule to the leaf tip on the adaxial surface of the first leaf above the top ear of the plant. Leaf width was measured on the same leaf at the midpoint between the ligule and the leaf tip. Extant leaf number was determined by counting the number of visible leaf collars on the same two plants. Plant height was measured between the soil surface and the flag leaf collar using a marked pole. Leaf Area Index values were estimated using a LAI-2200C Plant Canopy analyzer (LI-COR, Inc). For each plot one above canopy and three below canopy measurements were collected using the LAI-2200C’s 270-degree view cap. The three below canopy measurements collected diagonally in the space between the two rows of the plot. The first measure adjacent to one row, the second equidistant between the rows and the third adjacent to the second row. Leaf Area Index measurements were collected between July 28^th^ - August 12^th^, 2020.

All ears were harvested from eight semi-randomly selected plants per plot with edge plants being excluded when possible. Ear length, ear width, length of fill, kernel row number and number of kernels per row were hand measured or hand counted for six ears per plot or all ears when less than six ears were present (Figure S2). The average of individual ears were used to calculate plot level values. All harvested ears were weighed, shelled, and the resulting pooled grain was also weighed. Cob weight was calculated as the difference between ear weight and grain weight. Initial hundred kernel weight was calculated by counting and weighing 100 kernels per plot after shelling and pooling of grain. Grain moisture was measured using a Dickey-John GAC® 2500-AGRI Grain Analysis Computer (Dickey-John® Corporation, Auburn, IL). Total grain weight and hundred kernel weight was recalculated to a standardized 15.5% moisture content. When insufficient grain was harvested to collect accurate grain moisture data using the GAC® 2500-AGRI, a default value of 8.25% moisture, corresponding to the approximate median of all grain moisture values was employed to calculate moisture standardized grain weight and hundred kernel weight. Further, the BLUPs for each phenotype was calculated by fitting a linear mixed model using R package lme4^11^ with genotypes fit as random variable for the traits with data from 2020.

### Unified Genetic Marker Data

A single set of markers scored across 1,049 accessions were employed for downstream analyses. These genotypes were determined based on marker data aggregated from three published sources: resequencing data for the WiDiv-503 panel (454 individuals)^12^, resequencing data generated as part of the HapMap3 project (141 individuals)^7^, RNA-seq data for the WiDiv-942 panel (399 individuals)^5,6^. The specific NCBI SRA ID numbers of the files used to call SNPs for each of the accessions are provided in Table S2.

Both genome resequencing data and RNA-seq data were quality trimmed using Trimmomatic (v0.33)^13^. BWA-MEM (v0.7) with default parameter settings^14^ was employed to align the resulting trimmed resequencing data to v4 of the B73 maize reference genome^15,16^. STAR (v2.7)^17^ was used to align the trimmed RNA-seq reads to v4 of the B73 maize reference genome in two rounds as described in Sun et al^18^. Apparent PCR duplicates were marked within the resulting BAM alignments using picard (v2.22)^19^. *A priori* segregating genetic markers identified in maize HapMap3^7,9^ scored for each individual using GATK toolkit (v5.1)^20^. Missing values were imputed using beagle/5.01 with the HapMap3 population treated as a reference panel and parameters settings: ‘window=1 overlap=0.1 ne=1200’^21^. The imputed genetic marker dataset was filtered to remove markers with a minor allele frequency less than 0.01 or proportion of site heterozygous calls greater than 0.1 to produce the final set of 17,717,568 SNP markers.

### Quantitative Genetic Analysis of Trait Data

A kinship matrix for the complete set of 1,049 genotyped accessions, including all maize lines included in the SAM or WiDiv panels and 35 additional maize lines for which sequence data was generated and released as part of Mazaheri *et al.* 2019^6^ were calculated using the first method described by VanRaden (2008)^22^ as implemented in rMVP (v1.0.5)^23^. Narrow-sense heritability for each trait was calculated using this kinship matrix and the R package sommer (v4.1.1)^24^. Multidimensional scaling or the Principal Coordinate (PCo) Analysis was performed with –mds-plot 2 and –cluster options within plink v1.90^25^. Genome-wide patterns of linkage disequilibrium decay were estimated by calculating (LD/r2) for all pairs of genetic markers where both genetic markers exhibited minor allele frequency greater than 0.05 and were separated up to a physical distance of less than 600 kilobases using PopLDdecay (v3.41)^26^.

Marker-trait associations were identified using 100 iterations of the FarmCPU algorithm as implemented in the R package rMVP v1.0.5 with parameter settings maxLoop = 10; method.bin = “FaST-LMM”^23,27^. For each iteration, the first three principal components calculated from the genetic marker dataset were used as fixed effect and the kinship matrix calculated internally by the FarmCPU algorithm was fitted as random effects. The overall marker file was filtered on a per-GWAS basis to retain only those markers with a minor allele frequency >0.05 among the lines phenotyped a given trait prior to association testing for that trait. For each trait, 100 analyses were run, each incorporating data from a different randomly selected subset of phenotyped lines^28^. In each resampling analysis, the overall threshold for statistical significance was the bonferroni corrected p-value at 5%. Resampling model inclusion probability (RMIP) values for each marker were calculated as the proportion of the 100 analyses in which that marker was significantly associated with the target trait^28–32^.

Linkage disequilibrium was calculated among all genetic markers with RMIP values ≥5 for at least one trait. Genetic markers with linkage disequilibrium >0.5 were merged into single peaks for downstream analyses, unless the markers were separated by >1 Mb. When two or more markers were merged into a single peak, the marker with the greatest RMIP value was selected as representative of the entire peak.

### Data Availability Statement

Phenotypic values for all trait datasets employed in this study for all maize accessions evaluated are provided in Table S2 and S3. The sources of sequence data used to call genetic marker genotypes for each maize accession were NCBI BioProjects PRJNA661271, PRJNA189400 & PRJNA437324^5,6,8^. The specific SRA IDs for individual maize accessions are indicated in Table S2. The locations of GWAS peaks, the traits associated with each peak and the genes adjacent to each peak are provided in Table S4 and S5. The VCF file used for genetic analyses has been deposited at https://doi.org/10.6084/m9.figshare.19175888. Scripts and code used to implement various analyses described in the methods section above have been deposited in https://github.com/ravimural/Maize_resampling-based_GWAS.

## Results

### Properties of widely studied maize association panels

Three maize association panels were identified in the literature: the Maize Association Panel (MAP)^2^, the Shoot Apical Meristem (SAM) panel (369 lines)^3^ and the Wisconsin Diversity (WiDiv) panel consisting of either 627 or 942 lines^4,6^. The latter two populations are largely supersets of MAP, with SAM excluding 10 MAP lines present in the WiDiv panel and WiDiv excluding 67 lines retained in both the SAM and MAP populations (Figure 1A). The SAM and WiDiv populations also share 95 lines not present in the MAP population that served as a partial progenitor for both. These are predominantly more recently released lines developed in the private plant breeding sector and released when the associated plant variety parents expired (expired Plant Variety Patents or exPVPs). The total overlap between the SAM and WiDiv populations was 297, sufficient to enable joint analyses of trait datasets collected in these two populations. The union of the SAM and WiDiv populations included 1,014 unique maize genotypes with both genetic marker information at least one phenotypic record. An additional 35 unique maize genotypes where included in one or both populations, had at least one source of genetic marker data but no phenotypic records and so were excluded from downstream analyses.

**Figure 1.**
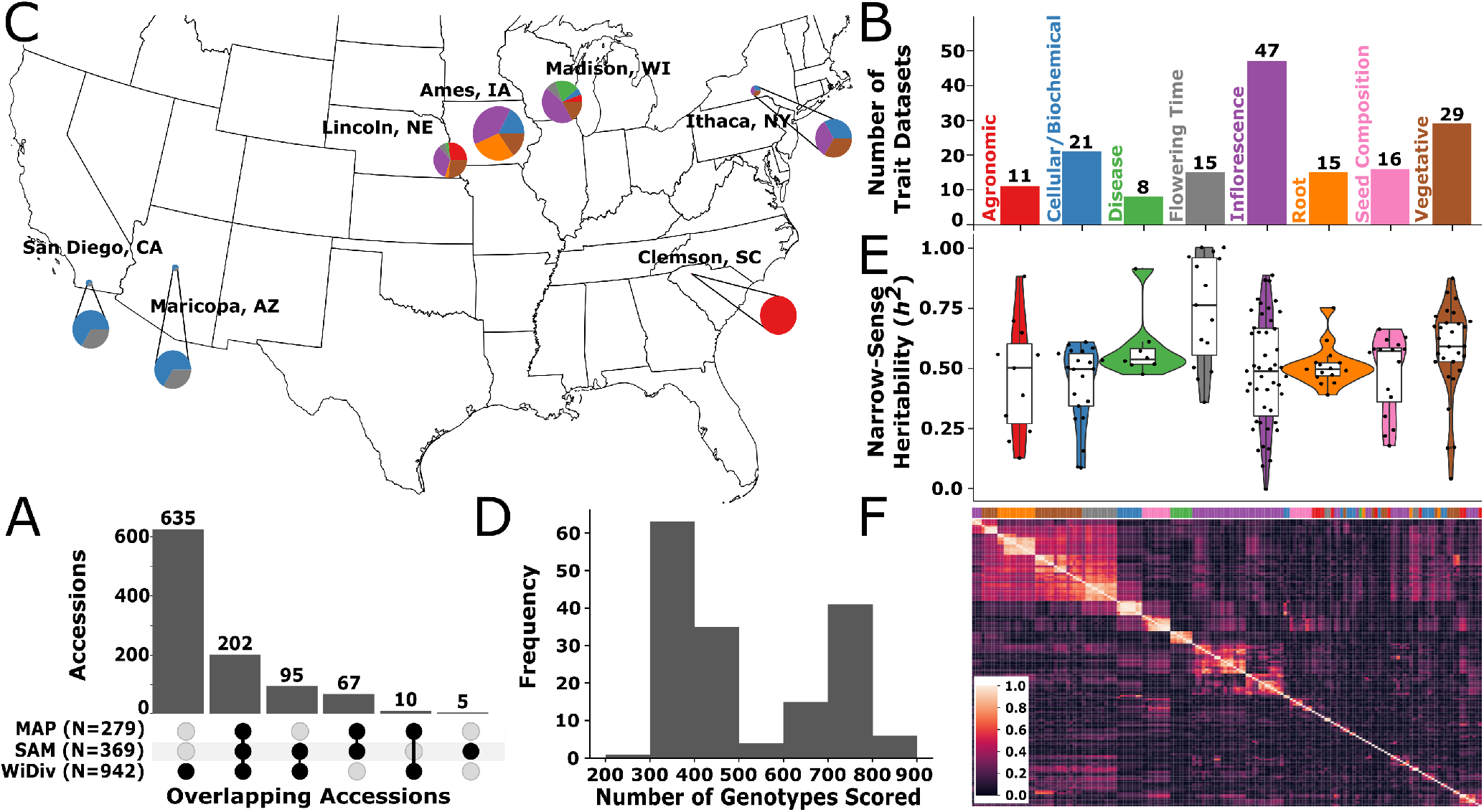
Characteristics of Maize Association Panel trait datasets. A) Number of accessions which are represented in any of the three diversity panels. B) Representation of eight broad phenotypic categories among the 162 traits collected here. Category assignments for individual traits are provided in Table S3. C) Geographic distribution of trials where trait datasets were collected. Size of circles indicates number of traits collected at a specific geographic location. Colors of circles indicate types of trait datasets collected at that location. Labels for which colors correspond to which types of traits are given in Panel B. D) Distribution of the number of genotypes scored for a given trait. E) Distributions of narrow-sense heritability values, across the same eight broad phenotypic categories shown in panel B. Colors corresponding to the color key for phenotype classes are provided in Panel B. F) Correlations among the 162 trait datasets analyzed in this study. Trait datasets are clustered based upon absolute spearman correlation value. Phenotype classes are indicated with color bar on top x-axis with colors corresponding to the color key for phenotype classes provided in Panel B.

The set of approximately 200 papers citing either the SAM panel^3^ or either iteration of the WiDiv panel^4,6^ were screened to identify studies which conducted GWAS and published trait datasets collected from one or both of these populations. A total of 21 papers were identified which included GWAS results generated from these populations. After excluding studies where we were unable to locate trait data for individual maize lines, excluding traits which failed initial QC, and condensing studies which utilized previously published data, 132 unique trait datasets drawn from 15 separate published studies remained. This included 55 trait datasets collected from the SAM panel, 66 trait datasets collected from the WiDiv panel, and 11 trait datasets collected from an even larger population of 2,815 maize lines with substantial overlap with these two populations^33^. An additional 30 phenotypes scored in Lincoln, Nebraska in 2020 were included for a final set of 162 trait datasets employed for downstream analyses (Table S3). Individual trait datasets included data values for between 222 and 817 maize lines (Figure 1B) and were collected from field or controlled environment studies conducted in seven states (Figure 1C). Measurements related to inflorescence architecture were the most abundant category among these 162 trait datasets (Figure 1D). SNP-based estimates of narrow-sense heritability for individual trait datasets were variable, with a median value of 0.527 and mean value of 0.523 across all traits. Traits related to flowering time (e.g. timing of anthesis, timing of silking, or anthesis-silking-interval) was the category which exhibited the highest median heritability of 0.762 (Figure 1E, and Table S3). Flowering time traits collected in different environments were correlated with each other and also exhibited notable correlations with a subset of both above-ground vegetative and below-ground root related traits (Figure 1F).

The total number of unique maize line names observed across these 162 trait datasets was 1,118 which was modestly more than the set of 1,014 unique genotypes present across the WiDiv and SAM mapping populations. We speculate that this difference may result from the inclusion of local checks or lines-of-interest or changes in naming convention or transcription errors which we were unable to resolve. For the 1,014 unique genotypes named as part of the SAM population^3^ or WiDiv population^6^, raw whole-genome sequencing or RNA-seq sequence data was aggregated from a number of sources (Table S2)^6,7,12^ as described in Sun *et al.* 2021^9^. Alignment of published sequence data from these sources to the maize reference genome (B73_RefGen_V4)^15,16^, scoring of *a priori* segregating SNPs from HapMap3^7^, imputation, and filtering resulted in a set of 17,717,568 with minor allele frequency >0.01 and heterozygosity rate of <0.1, leading to an average of one SNP per 120 bp (see Methods).

The 17,717,568 polymorphic markers chosen for downstream analysis were distributed roughly evenly across the ten chromosomes of maize, with local reductions in SNP density around centromeres/pericentromeric regions of each chromosome (Figure S3A). Rare SNPs with minor allele frequencies <0.1 were modestly more abundant than common SNPs (Figure 2A). Linkage disequilibrium decayed rapidly, with the average *r*^2^ between two SNPs separated by 10 kilobases being approximately 0.18 (Figure 2B) similar to previous reports^34,35^. The first three principal components of variation explained approximately 10% of total variance among genotypes (Figure S3B). Principal coordinate (PCo) analysis using this SNP set separated lines with known assignments to major heterotic groups (Figure 2C and D). The same set of PCo analyses did not identify obvious biases in the distribution of lines present in different association panels (Figure S3C).

**Figure 2.**
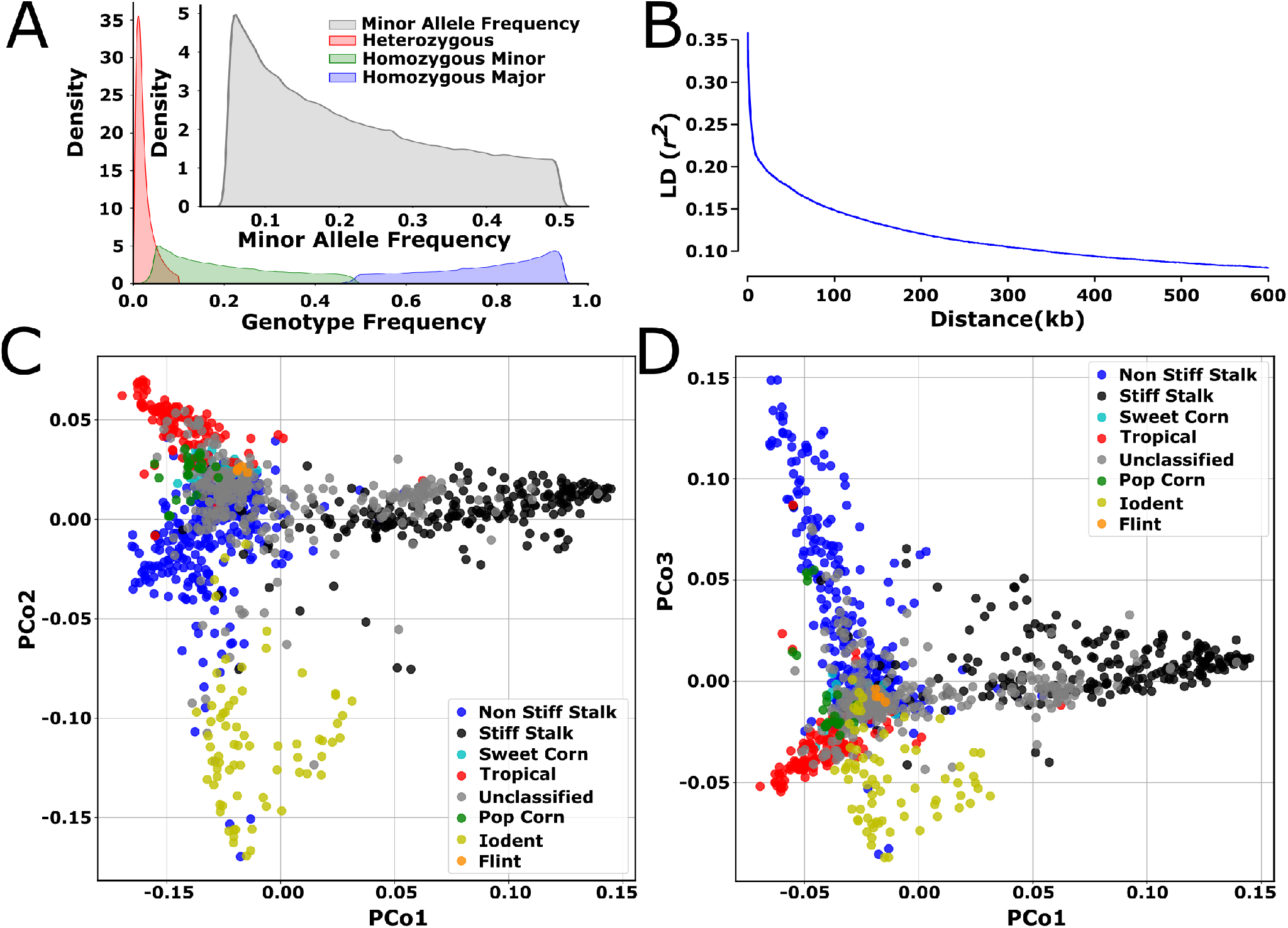
Characteristics of Maize Association Panel Marker datasets. A) Genotype frequency and minor allele frequency of the marker dataset. B) The genome-wide LD decay with maximum distance of 600 kilobases between two SNPs. C) Genetic relationship among the accessions used in this study and visualized using multidimensional scaling/principal coordinate analysis of the distance matrix. The X- and Y-axis represent first and the second principal component coordinates. Each point is color coded by the heterotic group each accession belongs to. D) Genetic relationship among the accessions used in this study and visualized using multidimensional scaling/principal coordinate analysis of the distance matrix. The X- and Y-axis represent first and the third principal component coordinates. Each point is color coded by the heterotic group each accession belongs to.

### Unified marker-trait analyses

Genome-wide association studies conducted using FarmCPU with the 162 traits and about 18M markers described in the previous section and employing a RMIP cutoff of 5 for a suggestive association identified 2,154 signals across 151 traits (Figure 3A). Among traits with one or more suggestively significant hits, the median number of hits was 12 (mean 12.57), the maximum was 33 and the minimum was 1.

**Figure 3.**
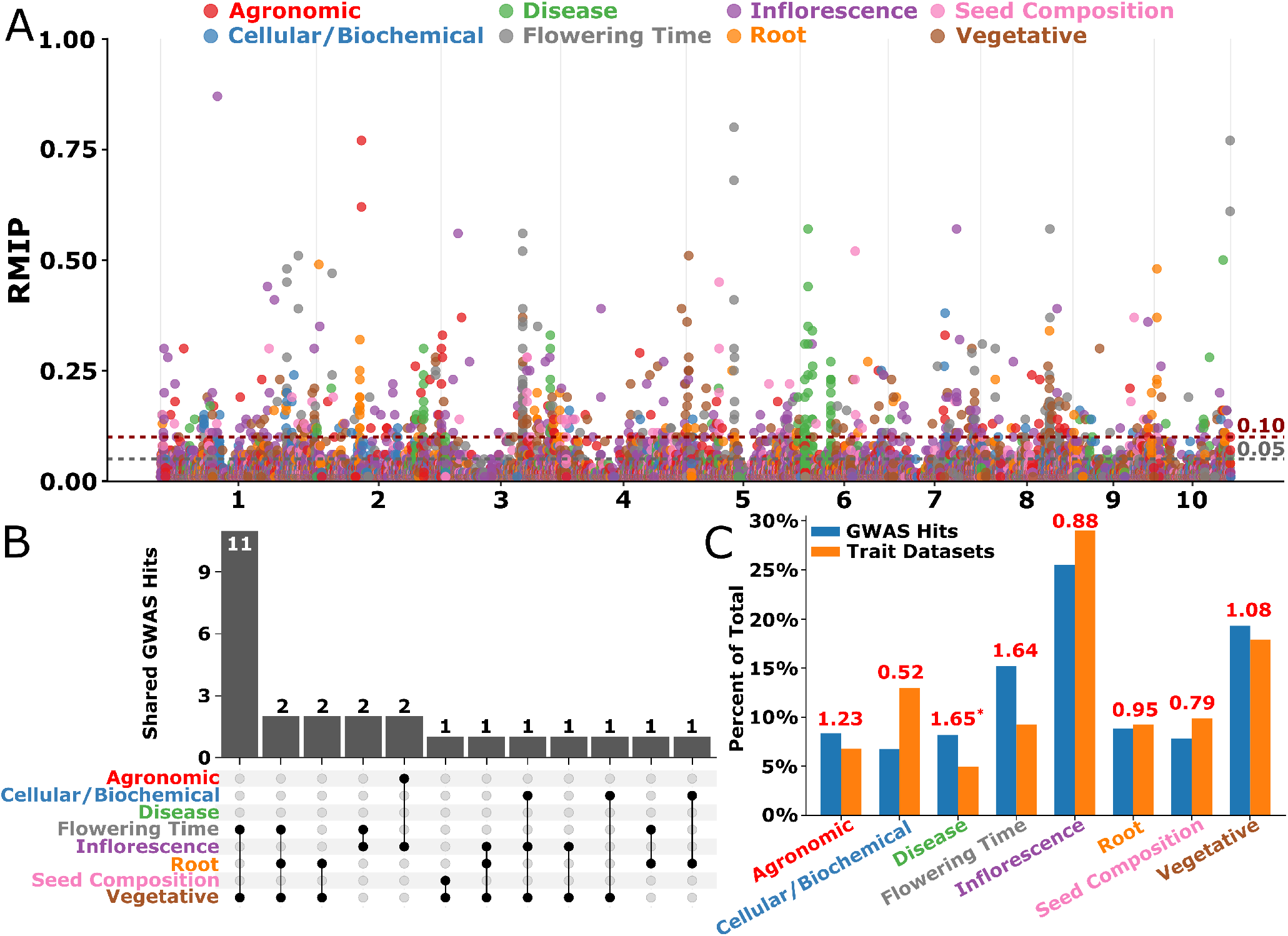
GWAS Summary: Multi-trait peaks detected across phenotypic categories. A) Combined Manhattan plot for GWAS using all 1014 individuals screened using 18M markers. The SNPs with RMIP ≥5 are plotted here. Dashed grey and red lines indicates the cutoff of 5% and 10% for statistical significance calculated based on RMIP value. Each chromosome is shown in the X-axis. The Y-axis is the resampling model inclusion probability (RMIP) values ranging from 0 to 1. B) An upset plot showing number of shared GWAS hits between various phenotypic categories. C) Percent representation of GWAS hits for the number of trait datasets analyzed. Number on top of each pair of bars in each phenotypic category corresponds to the ratio of GWAS hits:number of trait datasets analyzed in each category. *Note: The ratio was higher for the disease traits but the traits in this category are essentially the same trait analyzed at different timepoints in a time series manner, thus most of the hits overlap among the traits leading to an inflated ratio.

Consolidation of 2,154 SNPs with at least suggestive statistical significant associations with phenotypes (≥5 RMIP) into distinct peaks based on physical distance and LD (See Methods) reduced the number of associations to 1,466 peaks distributed across phenotypes assigned to eight categories (Table 2). Of these 1,466 peaks, 161 peaks were associated with 11 agronomic traits, 92 peaks were associated with 17 of the 21 total cellular/biochemical traits, 72 peaks are associated with eight disease traits, 176 peaks are associated with 15 flowering time traits, 459 peaks were associated with 41 of 47 total inflorescence traits, 113 peaks were associated with 15 root traits, 128 with 16 seed composition traits and 295 with 28 of 29 total vegetative traits (Figure 3B, and Table 2).

A wide range of approaches are employed in the literature to define the set of annotated gene models adjacent to a significant GWAS peak which should be labeled as “candidate genes”. These can include both fixed windows around the peak, examining an arbitrary number of the closest annotated gene models to the peak, or adaptive windows defined based on local levels of linkage disequilibrium or haplotype blocks. To assess the precision provided by the peaks identified in this study, we utilized a set of 604 gene models recorded in the MaizeGDB database^48^ as associated with one or more phenotypes. These gene models constituted 1.5% of the total set of 39,498 annotated gene models present on the B73_v4 reference genome^16^. The first three genes closest to GWAS peaks identified above were more likely to be associated with reports of phenotypes in MaizeGDB than the expected background rate, and this pattern became stronger at more stringent RMIP cutoffs (Figure 4A). When employing physical distance rather than rank order, the greatest enrichment of genes with reported phenotypes in the MaizeGDB database was observed in the categories “within gene” “closer than 10 kilobases” and “10-40 kilobases”, although noticeable enrichment remained observable at greater distances from the GWAS peak (Figure 4B).

**Figure 4.**
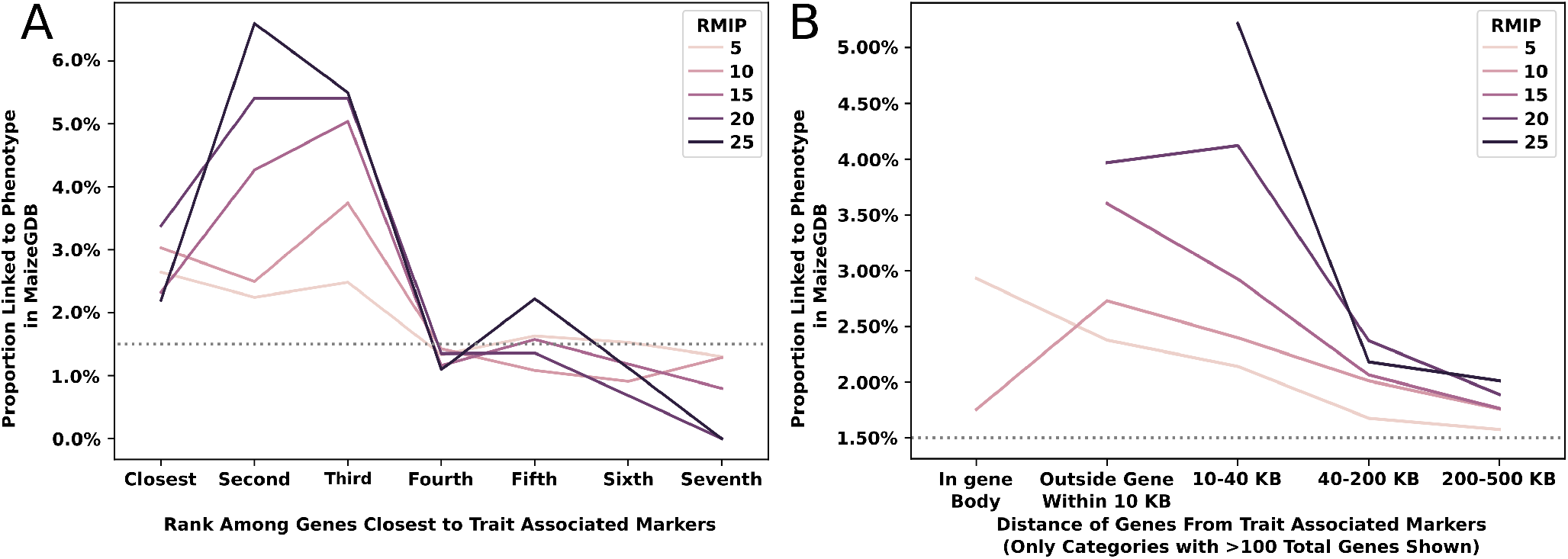
Probability of genes at different distances from peak SNP from GWAS are linked to phenotypes. A) Gene positions of unique trait associations. First seven genes closest to the GWAS peaks were selected and shown on X-axis. B) Gene order of unique trait associations. The distance of the genes from the trait associated markers are shown on X-axis.

Flowering time trait datasets tended to identify a disproportionately high number of independent GWAS peaks (Figure 3C), potentially as a result of the greater proportion of variance among these traits explained by genetic factors (Figure 1E). Overall, in 1,252 cases (85.4%) a peak was identified only in the analysis of a single trait dataset (Table 2). The remaining 214 peaks were identified in analyses of two or more separate trait datasets. In 188 cases the same peak was identified in the analysis of two or more phenotypes belonging to the same general category. For example, a peak consisting of four SNPs in high LD with each other on chromosome 6 spanning from 108,211,603 to 108,213,234 bp with the single highest RMIP SNP located at 108,212,338 bp was identified in analysis of both kernel starch abundance (Starch_K) (RMIP=52) and kernel fat abundance (Fat_K) (RMIP=23) within the overall category of “seed composition” traits^45^. The peak spans the 5’ end of the gene model Zm00001d036982 (108,212,462 to 108,219,350 on chromosome 6) which encodes *DGAT1-2* (Diacylglycerol O-acyltransferase 1-2)/ *ln1* (linoleic acid1). *DGAT1-2* substantially increases the seed oil and oleic-acid contents^49^. Largely, oils are stored in the form of triacylglycerol (TAG) and DGAT catalyzes the final step of TAG biosynthesis by transferring an acyl group from acyl-CoA to the sn-3 position of 1,2-diacylglycerol (DAG) thus, acting as the rate-limiting enzymes for TAG biosynthesis^50,51^ (Figure 5A&B). The rarer allele (“T”) is associated with an increase in seed fat and decreases in seed starch (Figure 5C). The starch-promoting allele was more abundant in iodent subpopulations and less abundant in sweet corn subpopulations (Figure S4). The original analysis of these two datasets employed the FarmCPU algorithm, but without resampling^45^ and did not identify these two associations, consistent with observed RMIP values which suggest only a one in two chance of detecting the DGAT/starch association and only a one in four chance of detecting the DGAT/fat association in a single round of GWAS.

**Figure 5.**
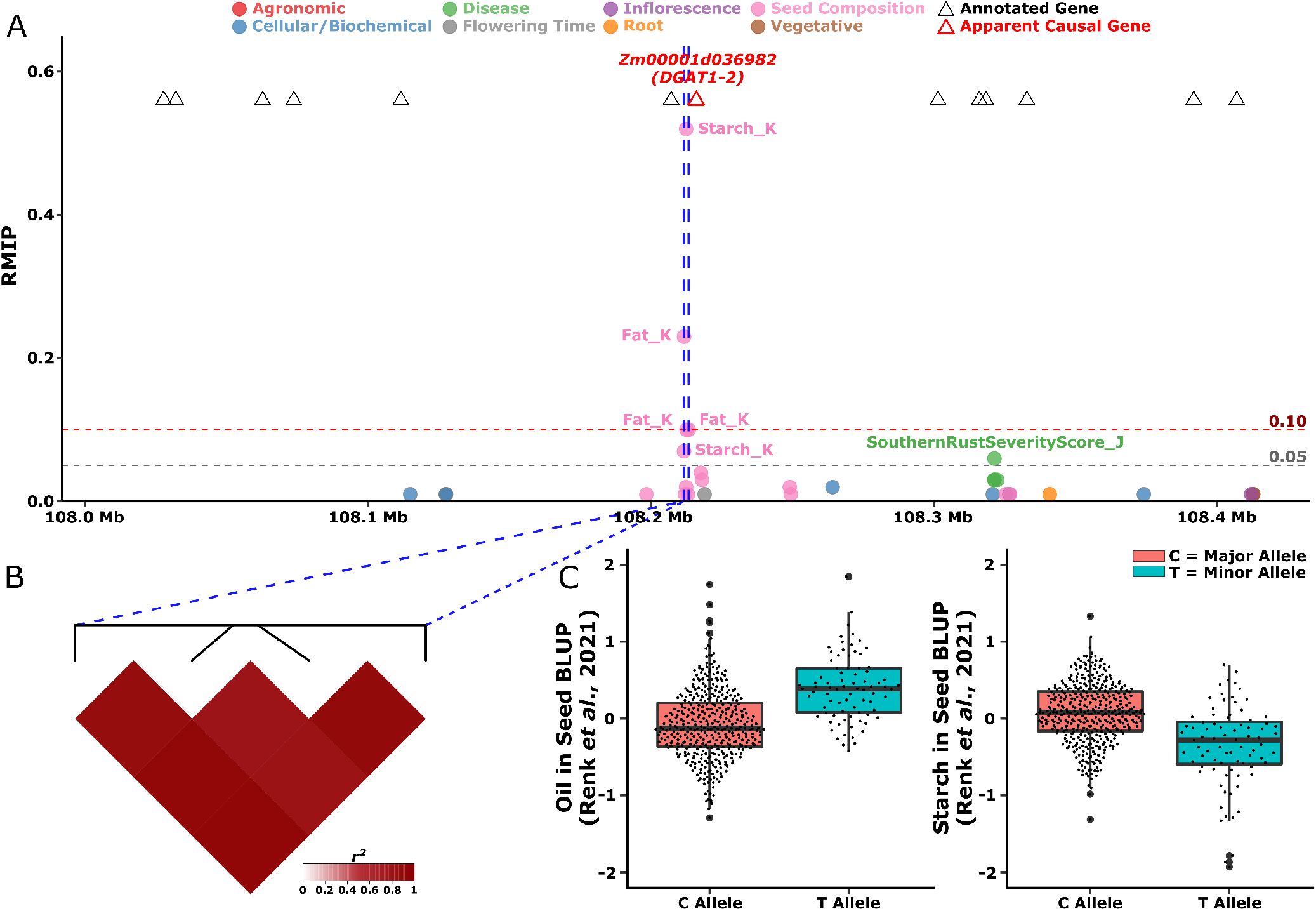
Combined GWAS identifies peak associated with seed starch and fat. A) View of resampling marker inclusion probability values for markers in a window from 108,211,603-108,213,234 on chromosome 6 spanning 200 kilobases upstream and downstream of the pleiotropic peak identified for seed starch and oil content. Only markers with resampling marker inclusion probability values ≥ 0.01 are shown. B) The LD relationships between the significant SNPs within the peak. C) Distributions of observed oil and starch content values reported in^45^ for lines carrying either allele of the peak SNP located at position 108,212,338 bp.

In the remaining 26 cases where the same genomic interval was identified in the analysis of multiple trait datasets (Figure 6A–D, and S5–S26), the trait datasets involved spanned two or more categories, with 22 peaks associated with trait datasets spanning two categories and four peaks associated with trait datasets spanning three categories (Figure 3B). Genomic intervals associated with flowering time were disproportionately more likely to be associated with phenotypes from at least one other category. Sixteen of 176 unique peaks identified for flowering time were also associated with one or more phenotypes from other categories (9%), while only 10 of 1,290 unique peaks (0.8%) identified for non-flowering time traits were associated with traits from two or more of the remaining seven categories (Figure 3B).

**Figure 6.**
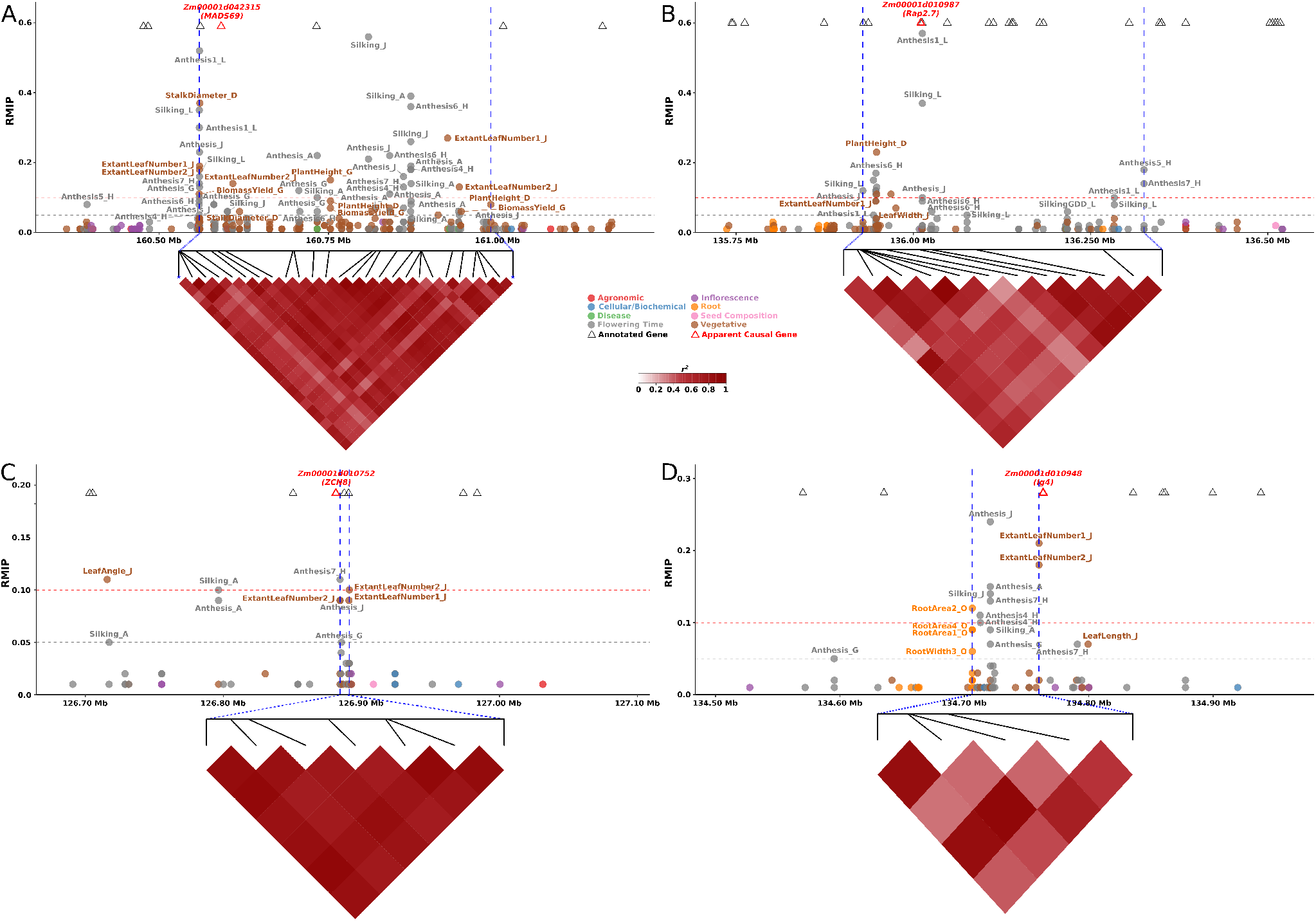
GWAS peaks associated with multiple traits. A) Local Manhattan plot with + /−200 kilobases of pleiotropic peak on chromosome 3 from 160,559,294 to 160,989,691 bp. This peak is associated with *MADS69* (Zm00001d042315). The phenotypes associated with this peak belongs to Flowering Time and Vegetative categories. The phenotypes associated with this peak are Anthesis1_L, Anthesis4_H, Anthesis6_H, Anthesis7_H, Anthesis_A, Anthesis_G, Anthesis_J, BiomassYield_G, ExtantLeafNumber1_J, ExtantLeafNumber2_J, PlantHeight_D, PlantHeight_G, Silking_A, Silking_J, Silking_L, and StalkDiameter_D. The vertical dashed lines shows the peak boundary. B) Local Manhattan plot with + /−200 kilobases of pleiotropic peak on chromosome 8 from 135,928,821 to 136,325,345 bp. This peak is associated with *Rap2.7* (Zm00001d010987). The phenotypes associated with this peak belongs to Flowering Time and Vegetative categories. The phenotypes associated with this peak are Anthesis1_L, Anthesis5_H, Anthesis6_H, Anthesis7_H, Anthesis_A, Anthesis_G, Anthesis_J, ExtantLeafNumber1_J, LeafWidth_J, PlantHeight_D, SilkingGDD_L, Silking_L. The vertical dashed lines shows the peak boundary. C) Local Manhattan plot with + /−200 kilobases of pleiotropic peak on chromosome 8 from 126,884,534 to 126,891,234 bp. This peak is associated with *ZCN8* (Zm00001d010752). The phenotypes associated with this peak belongs to Flowering Time and Vegetative categories. The phenotypes associated with this peak are Anthesis7_H, Anthesis_G, Anthesis_J, ExtantLeafNumber1_J, ExtantLeafNumber2_J. The vertical dashed lines shows the peak boundary. D) Local Manhattan plot with + /−200 kilobasesof pleiotropic peak on chromosome 8 from 126,884,534 to 126,891,234 bp. This peak is associated with *lg4* (Zm00001d010948). The phenotypes associated with this peak belongs to Flowering Time, Root and Vegetative categories. The phenotypes associated with this peak are Anthesis4_H, Anthesis7_H, Anthesis_A, Anthesis_G, Anthesis_J, ExtantLeafNumber1_J, ExtantLeafNumber2_J, RootArea1_O, RootArea2_O, RootArea4_O, RootWidth3_O, Silking_A, Silking_J. The vertical dashed lines shows the peak boundary.

An illustrative example of the potential for genes influencing flowering time to be identified in genome-wide association studies for other traits is the case of *ZmMADS69*. *ZmMADS69* (syn *Zmm22*) (Zm00001d042315) is a MADS-box transcription factor located between 160,564,021 bp and 160,591,933 bp on maize chromosome 3, which has been shown to function as flowering activator, with a derived allele conferring earlier flowering in many maize lines relative to its wild progenitor teosinte^6,52^. A peak consisting of 26 SNPs in high LD with each other was consistently identified for multiple flowering time related traits including seven measurements of anthesis (male flowering) in different environments (Anthesis_A, Anthesis_G Anthesis1_L, Anthesis4_H, Anthesis6_H, Anthesis7_H, Anthesis_J) and three measures of silking in different environments (Silking_A, Silking_L, Silking_J). The same peak was also identified in the analysis of multiple vegetative traits including measurements of plant height in two environments (PlantHeight_D, PlantHeight_G), extant leaf number, stalk diameter, and biomass yield (Figure 6A). *ZmMADS69* has been shown to downregulate the expression of *ZmRap2.7*, which relieves repression of the florigen gene *ZCN8*, causing/resulting in early flowering^52^. Both *ZmRap2.7* and *ZCN8* are located on chromosome 8^53–55^ and both of these genes are also associated with GWAS peaks. A peak on chromosome 8 consisting of four SNPs between 126,884,534 bp to 126,891,234 bp was separated by only two kilobases from the gene model encoding *ZCN8* (Zm00001d010752, located between 126,880,531 and 126,882,389 bp) and was associated with anthesis in three environments (Anthesis_G, Anthesis7_H, Anthesis_J). *ZmRap2.7* (Zm00001d010987) is located approximately 10 megabases away from *ZCN8* on chromosome 8 (between 136,009,216 and 136,012,084 bp) and is associated with a peak consisting of 13 SNPs that was detected for seven measurements of anthesis (Anthesis_A, Anthesis_G, Anthesis1_L, Anthesis5_H, Anthesis6_H, Anthesis7_H, Anthesis_J), two approaches to measuring silking in the same environment (Silking_L, and SilkingGDD_L), and a number of vegetative traits including extant leaf number (ExtantLeafNumber1_J), leaf width (LeafWidth_J), and plant height (PlantHeight_D).

In addition to the three peaks discussed above, thirteen other peaks were also associated with both flowering time traits and traits from other categories (Figure 3.A, Figure 3.B, and Table S4). In two cases a significant signal for flowering time was co-located with significant signals for inflorescence architecture traits. The first, located on chromosome 1 between 102,077,749 and 102,120,437 bp, consists of two SNPs in high LD with each other and is significantly associated with both date of silking (Silking_J) and ear length (EarLength_O) in separate environments (Table S4, and Figure S5). The second, located on chromosome 4 between 78,020,118 and 78,451,569 bp, consists of two SNPs and showed significant associations with both male flowering in one environment (Anthesis5_H) and the length of the central spike of the tassel in another environment (SpikeLength1_C) (Table S4, and Figure S14). In eleven cases a significant association for flowering time was co-located in the genome with a signal for an above-ground vegetative trait dataset. These were typically vegetative traits with known links to flowering time including leaf/node number, and plant or ear height (Table S4, and Figure 6D, S6, S12, S16 - S20, S22, S24, and S26).

The potential pleiotropy of genes linked to flowering time was not confined to above-ground traits. In three cases a significant signal for flowering time was also associated with one or more datasets describing variation in root phenotypes. A signal on chromosome 5 between 94,710,702 and 94,712,951 bp was associated with flowering time across a wide range of environments (Anthesis_A, Anthesis1_L, Anthesis7_H, Anthesis_J, Silking_J, Silking_L, SilkingGDD_L), with other above-ground vegetative traits (LeafAreaIndex_J, LeafLength_J) and with many root architecture traits (RootArea1_O, RootArea2_O, RootArea4_O, RootWidth4_O) (Table S4 and Figure S19). The specific SNPs which define the peak are all located within Zm00001d015513 which encodes a cinnamoyl-CoA reductase expressed primarily in leaves and leaf meristems^56^. A signal on chromosome 8 between 28,727,658 and 28,769,198 bp, was associated with both male and female flowering time in Nebraska (Anthesis_J, Silking_J) and variation in root depth in Iowa (RootDepth1_O, RootDepth2_O) (Table S4 and Figure S22). The last of the three signals, associated with flowering time (Anthesis_A, Anthesis_G, Anthesis4_H, Anthesis7_H, Anthesis_J, Silking_A, Silking_J), leaf number, and root (RootArea1_O, RootArea2_O, RootArea4_O, RootWidth3_O) traits is also located on chromosome 8, between 134,706,389 to 134,759,977 bp. This 54 kilobase interval is entirely free of annotated genes, but ends 600 bp upstream of classical mutant *liguleless4* (Zm00001d010948) (Figure 6D). *Liguleless4* (synonym *knox11*) encodes a knox transcription factor which is highly expressed in the SAM, seed radicle, internode tissues, crown roots, pericarp of seed and the endosperm of maize^56^. A dominant allele of *liguleless4* abolishes the ligule and alters the sheath-blade boundary in maize leaves^57^ likely via ectopic expression^58^, however phenotype of loss of function alleles, if any, remains uncharacterized.

## Discussion

The widespread adoption of diverse association panels in plant biology has enabled a wide range of research and discovery by researchers working on diverse phenotypes, species, and research questions^59^. Beyond lists of specific candidate genes identified in the main text or supplemental figures, the reuse of GWAS results can sometimes be challenging. Changes in genome versions, gene model annotations, or genetic marker datasets, as well as changes in best practices and algorithms for conducting genome-wide association tests can all hinder comparisons with and/or reuse of previously published GWAS results. In the field of human genetics where the release of individual-level trait and genetic data could raise privacy and ethics concerns, the field has converged on a standard of releasing summary statistics but not individual-level trait and genetic data^60^. Working with plant data, privacy concerns do not typically preclude the release of individual-level data, and the reuse of the same genotypes across independent studies increases the potential value of individual-level data.

We identified 21 published papers describing phenotypes or GWAS conducted using one or more of two widely adopted association panels in maize^3,4^. In 18 cases (85%) it was possible to locate raw trait values for individual lines used in the published analyses, including the 16 studies summarized in Table 1 and the two additional papers not included in our analyses. The high frequency with which trait data is being released for published GWAS studies is encouraging as it indicates the maize quantitative genetics community is adopting similar norms and practices to the genomics community which has long been a leader in promoting strong best practices for raw data deposition and dissemination^61^. Unlike the genomics community, the plant quantitative genetics community does not have access to widely used and standardized data repositories. Challenges in integration of these data included inconsistent naming, extra lines that were not part of panel, repeated traits across papers, and data distributed across supplemental files or Figshare. Metadata for how and when individual traits were collected typically was present but often needed to be manually extracted from reading the manuscript text. Information that would further increase the value of released trait data such as the GPS coordinates and planting dates of individual field trials was provided in some cases but not others. The identification of a single common repository, standards for metadata on individual field trials, and for the preservation of a single unique identifier for each genotype included in a community association population would all lower barriers to the reuse of trait datasets. However, despite these current challenges both the overall consistency and quality of data release and documentation was exceptional, enabling the investigation of multi-environment and multi-trait genetic associations.

**Table 1.** Studies from which maize trait datasets were drawn

| Reference | Study Type | Phenotypes Scored<sup>a</sup> | Accessions Evaluated<sup>b</sup> | Panel |
| --- | --- | --- | --- | --- |
| Peiffer *et al.* 2014<sup>36</sup> | Reproductive & Vegetative | 11 | 737 | Ames Panel |
| Hirsch *et al.* 2014<sup>5</sup> | Reproductive & Vegetative | 3 | 427 | WiDiv-503 |
| Leiboff *et al.* 2015<sup>3</sup> | Agronomic, Cellular/Biochemical, & Vegetative | 9 | 378 | SAM |
| Lin *et al.* 2017<sup>37</sup> | Cellular/Biochemical, Root, & Vegetative | 16 | 363 | SAM |
| Gustafson *et al.* 2018<sup>38</sup> | Disease | 7 | 447 | WiDiv-503 |
| Gage *et al.* 2018<sup>39</sup> | Reproductive | 16 | 817 | WiDiv-942 |
| Mazaheri *et al.* 2019<sup>6</sup> | Cellular/Biochemical & Vegetative | 5 | 788 | WiDiv-942 |
| Qiao *et al.* 2019<sup>40</sup> | Cellular/Biochemical | 4 | 429 | WiDiv-503 |
| Sekhon *et al.* 2019<sup>41</sup> | Agronomic | 3 | 364 | WiDiv-503 |
| Zheng *et al.* 2019<sup>42</sup> | Agronomic, Root | 13 | 359 | SAM |
| Azodi *et al.* 2020<sup>43</sup> | Reproductive & Vegetative | 3 | 388 | WiDiv-503 |
| Lin *et al.* 2020<sup>44</sup> | Cellular/Biochemical & Reproductive | 8 | 439 | WiDiv-503 |
| Renk *et al.* 2021<sup>45</sup> | Seed Composition | 16 | 499 | WiDiv-503 |
| Schneider *et al.* 2021<sup>46</sup> | Root | 1 | 599 | WiDiv-503 |
| Zhou *et al.* 2021<sup>47</sup> | Reproductive | 17 | 339 | SAM |
| Sun *et al.* 2021<sup>9</sup> | Disease | 1 | 687 | WiDiv-942 |
| Previously Unpublished | Agronomic, Disease, Reproductive, & Vegetative | 29 | 752 | WiDiv-942 |

\*Note: "WiDiv": Wisconsin Diversity Panel, "SAM": Shoot Apical Meristem Panel, "UNL": University of Nebraska-Lincoln, "a": Phenotypes used in this study from the phenotypes scored in respective studies (after removing exact same phenotype values if used in another study).
\*Note: The highest number of accessions with phenotype data used in this study from the respective publication

**Table 2.** Summary of unique associations with RMIP ≥ 5 within each of the eight phenotypic groups analyzed.

| Phenotype Group | # of Phenotypes analyzed | # of Phenotypes with hits | # of peaks | # of single trait peaks | # of Multi Trait Peaks | # of Multi Trait Peaks within each Category <sup>a</sup> | # of Peaks Associated Across Category <sup>b</sup> |
| --- | --- | --- | --- | --- | --- | --- | --- |
| Agronomic | 11 | 11 | 161 | 155 | 6 | 4 | 2 |
| Cellular/Biochemical | 21 | 17 | 92 | 69 | 23 | 20 | 3 |
| Disease | 8 | 8 | 72 | 44 | 28 | 28 | 0 |
| Flowering Time | 15 | 15 | 176 | 128 | 48 | 32 | 16 |
| Inflorescence | 47 | 41 | 459 | 420 | 39 | 32 | 7 |
| Root | 15 | 15 | 113 | 81 | 32 | 25 | 7 |
| Seed Composition | 16 | 16 | 128 | 108 | 20 | 19 | 1 |
| Vegetative | 29 | 28 | 295 | 247 | 48 | 28 | 20 |
| Total Unique | 162 | 151 | 1466* | 1252 | 214* | 188 | 26* |

<sup>a</sup> Excluding 26 peaks that overlap between two or more phenotype groups/categories. Of these 26 peaks 22 peaks are associated with traits belonging to two phenotype categories and 4 peaks are associated with phenotype traits belonging to three phenotype categories.
\* The value is less than the total of all the values in respective columns because of peaks associated with phenotypes in multiple categories are depicted in each category they show significance.

One desirable outcome of having access to raw trait data is that it enables reanalysis of existing trait datasets, each of which represents a substantial investment of both finance resources and human effort/suffering, as new higher resolution genetic marker datasets and new analysis algorithms become available. Here we employed a RMIP-based filter to the FarmCPU GWAS algorithm^27–29^ and were able to identify 2,154 suggestive associations (RMIP≥5) and 697 confident associations (RMIP≥10) across 162 traits collected in 33 environments spanning at least seven states. These signals included new associations identified as a result of either new genetic marker data and/or the FarmCPU/RMIP approach (Figure 5). Overall 1,466 and 468 unique sites in the genome tagged with a suggestive or confident association, respectively. These associations were enriched near genes with previously reported phenotypic effects (Figure 4). However, many of these signals are located near genes whose functions were previously entirely unknown or estimated purely via functional data on homologs which is typically useful for inferring molecular function of a protein encoded by a given gene but can produce misleading information on the specific biological processes a given gene is involved in^62^. These new layers of functional data will be most useful if they can be integrated into community genomics repositories. In this case the functional data generated as part of this project have been integrated as browser tracks and downloads at the maize community repository MaizeGDB^48^ to enable maize researchers to quickly access these data, cross reference them with other data types, and compare with mutant or QTL mapping results.

The significant signals identified for flowering time and both above-ground and below-ground plant architectural traits adjacent to *liguleless4*/*knox11* are an example of an intermediate case between confirmation of known functions and assigning potential functions to previously uncharacterized genes or regions of the genome. *Liguleless4* belongs to the knox class-I gene family^58^ a family of genes involved in regulating plant development via expression in apical meristems^63^. The *liguleless4* gene itself was identified via a dominant allele *Lg4-O* which alters development at the leaf blade/sheath boundary^57^ and associated with ectopic expression^58^. Loss of function alleles of the *liguleless3* gene, a paralog of *liguleless4*, do not exhibit any obvious phenotype^58^. A role for *liguleless4* in determining flowering time and above/below-ground plant architecture is consistent with the reported expression pattern of the wild type allele in tissues including the shoot apex, root tips, and developing inflorescence^58^.

One striking observation from the colocalization of association signals across multiple trait datasets was how common the re-identification of shared signals was. 14.5% (214/1466) of all suggestive associations and 16.2% (76/468) of all confident associations were identified in at least two trait datasets. One utility of reanalyzing published trait datasets is that variants with consistent but moderate effects across many studies can be distinguished from, and assigned higher confidence, than signals of equivalent statistical significance which are identified in only a single study in a single environment. Another lesson to take away from the same colocalization data is how common it was for the same locus to be identified for traits belonging to separate categories of phenotypes. The interpretation of a genetic locus with a significant association with root area will be quite different depending on whether than same locus is also associated with flowering time^64^ (Figure 6D, S19 and S22). In both cases the key take away is that researchers do not have to analyze or interpret GWAS in a vacuum but instead are able to interpret their results in the context of the rich datasets of previously scored phenotypes and previous GWAS analyses. Our understandings of genetics, genotype by environment interactions, and pleiotropy will all benefit from the broad use of these rich datasets.

## Acknowledgements

The authors thank Thomas Hoban, Isabel Sigmon, Alice Guo, Nathaniel Pester, Leighton Wheeler, Sierra Conway, Isaac Stevens, and Olivier N. Mizero for assistance with field maintenance, harvest, and trait data collection. This material is based upon work supported by the US Department of Energy Advanced Research Projects Agency-Energy (ARPA-E) under Award Nos. DE-AR0001064 and DE-AR0001367, the National Science Foundation under grants OIA-1557417 and OIA-1826781 and USDA-NIFA under the AI Institute: for Resilient Agriculture, Award No. 2021-67021-35329 and the Foundation for Food and Agriculture Research Award No. 602757. This research was supported in part by the US. Department of Agriculture, Agricultural Research Service Project #5030-21000-068-00D. Mention of trade names or commercial products in this publication is solely for the purpose of providing specific information and does not imply recommendation or endorsement by the U.S. Department of Agriculture. USDA is an equal opportunity provider and employer.

## Author contributions statement

RVM, and JCS conceived of the study. RVM, GS, MG, MCT, HJ, CS, LN, AMT, and BS designed and carried out experiments to generate data. RVM, MG, and GS analyzed the data. RVM and JCS wrote the initial draft of the manuscript. All authors read and approved the final manuscript.

---

**Figure S1.**
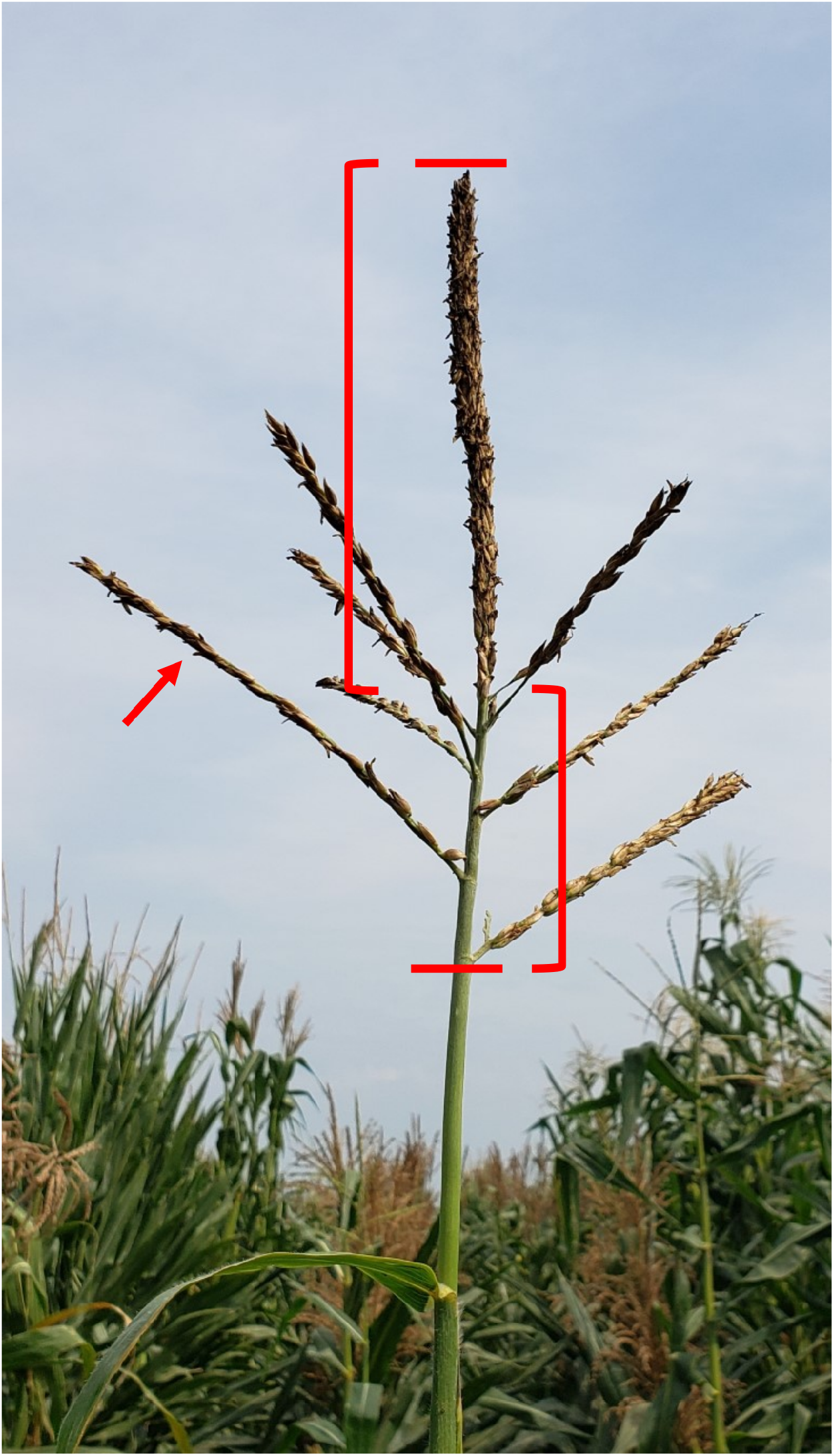
Phenotyping of Tassel Architecture. A) Tassel lengths were measured from the bottom-most primary tassel branch to the tip of the tassel spike (red bars). B) The branch zone length was defined as the length from the bottom-most primary tassel branch to the upper-most primary tassel branch (red left-facing bracket). C) The tassel spike length was defined as the length from the upper-most primary tassel branch to the tip of the tassel spike (red right-facing bracket). D) The total number of primary tassel branches were also counted (example arrowed) as well as the number of these primary tassel branches that were initiated, but later aborted (not pictured).

**Figure S2.**
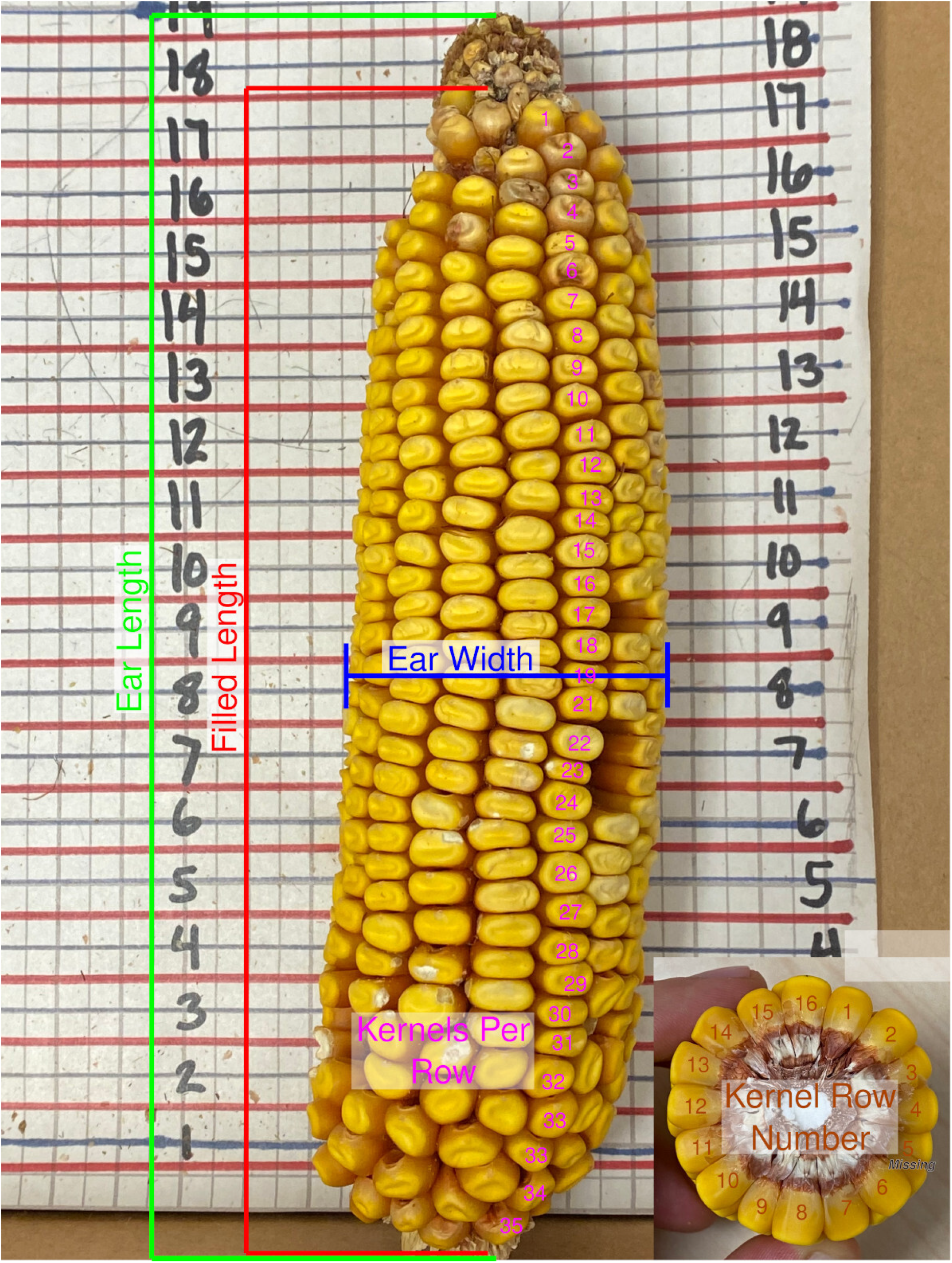
Phenotyping of Cob Traits. A) Ear lengths were measured as distance from base to the tip of each cob in centimeters, highlighted in green. B) ear fill was measured as distance on the cob from base to the tip where the seeds were set, highlighted in red. C) Ear width was measured as distance of diagonal of the cob, highlighted by blue color D) Kernels per row corresponds to the number of kernels on each line when cobs were set vertically, highlighted in pink color. E) Kernel row number correspond to the number of rows, highlighted in Brown color.

**Figure S3.**
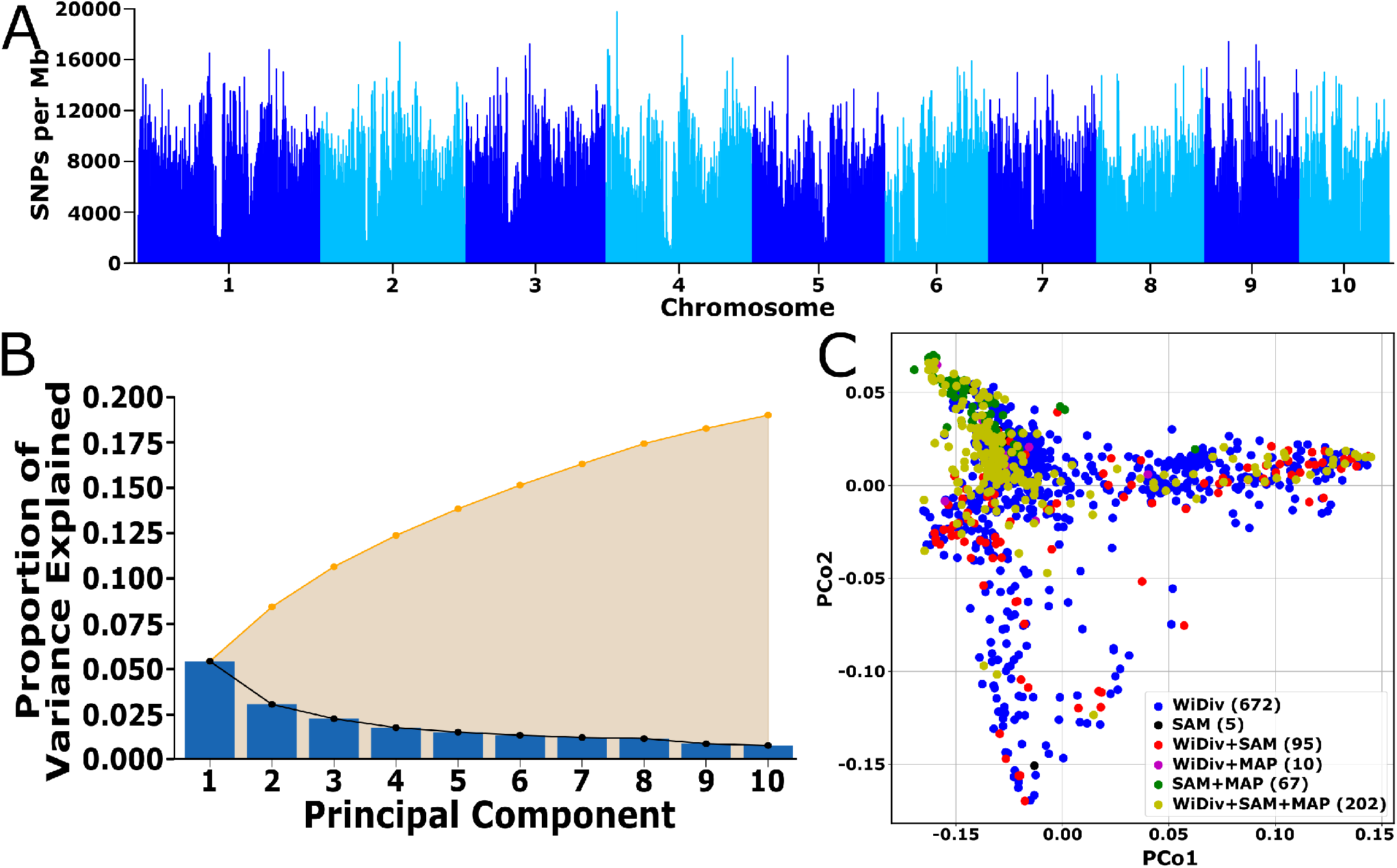
Multi Dimension Scaling or Principal Coordinate Analysis. A) Distribution of SNP density across the sorghum genome in 1 megabase sliding windows. B) Scree plot of eigen values for the principal components estimated from the marker data used in this study. C) Genetic relationship among the accessions used in this study and visualized using multidimensional scaling/principal coordinate analysis of the distance matrix. The X- and Y-axis represent first and the second principal component coordinates. Each point is color coded by the community association panels each accession belongs to.

**Figure S4.**
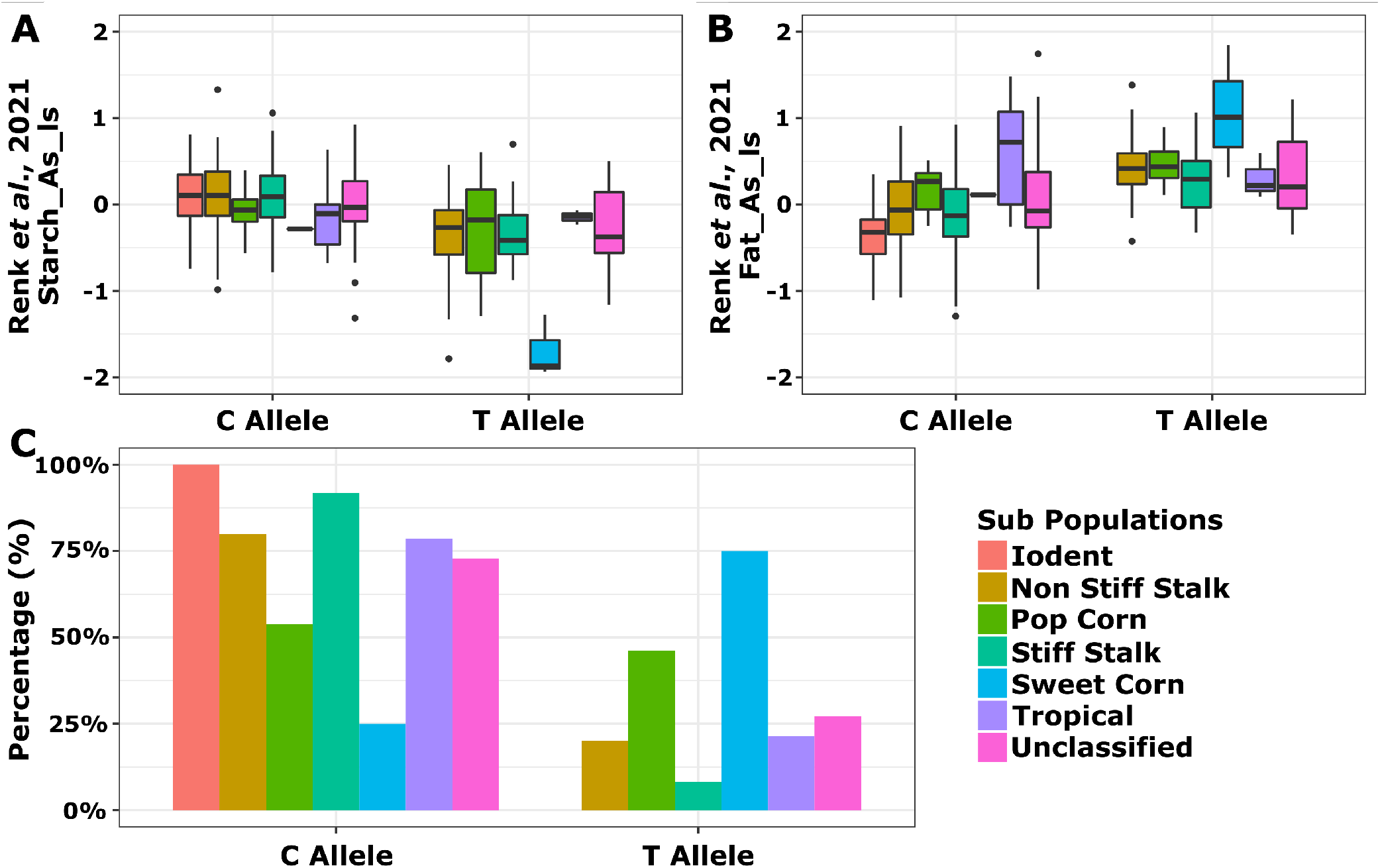
Allele frequencies of the top SNP associated with the DGAT-2 gene. A) The allele frequency for the starch content in each subpopulation of top SNP associated with DGAT gene. B) The allele frequency for the fat content in each subpopulation of top SNP associated with DGAT gene. C) Percentage of alleles in each subpopulation: The starch promoting allele was more abundant in iodent subpopulations and less abundant in sweet corn subpopulations.

**Figure S5.**
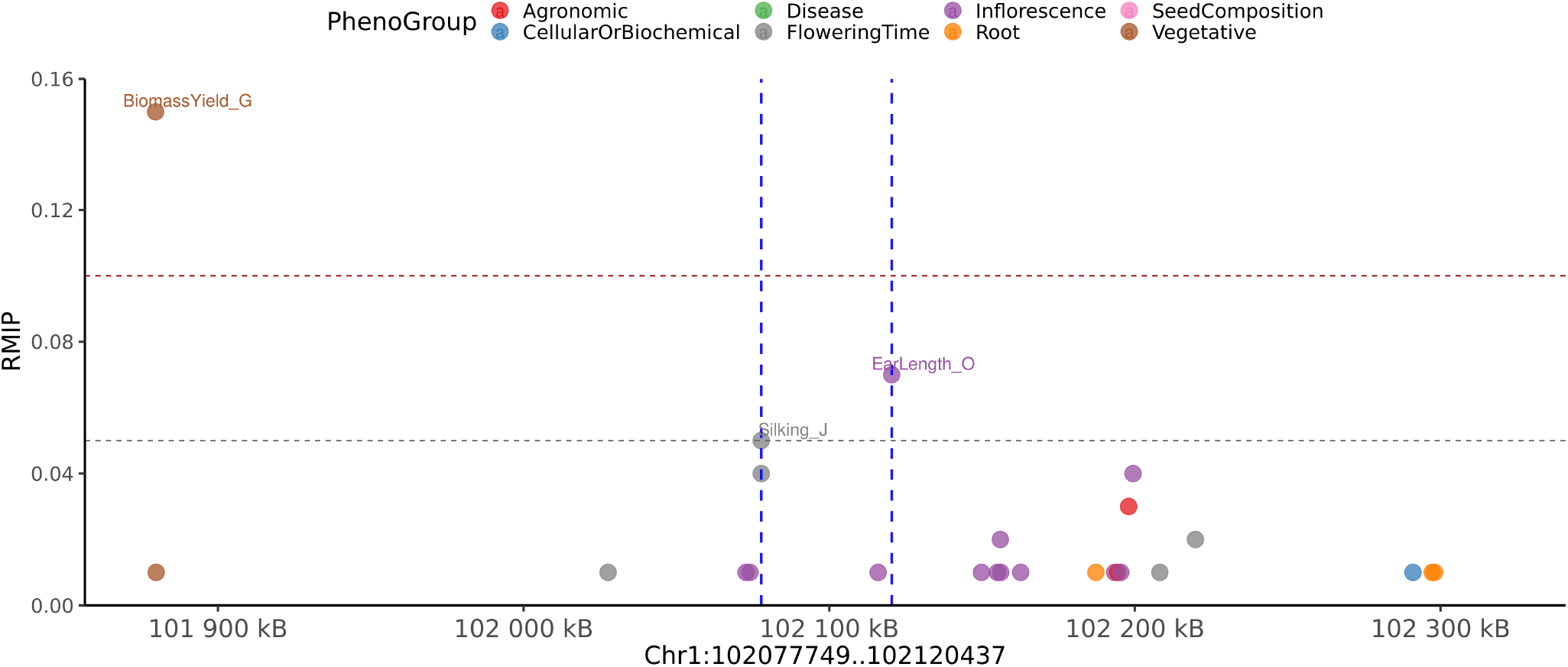
Local Manhattan plot with + /−200 kilobases of pleiotropic peak on chromosome 1 from 102,077,749 bp and 102,120,437. This peak is associated with the phenotypes belonging to Inflorescence and Flowering Time categories. The phenotypes associated with this group are EarLength_O and Silking_J.

**Figure S6.**
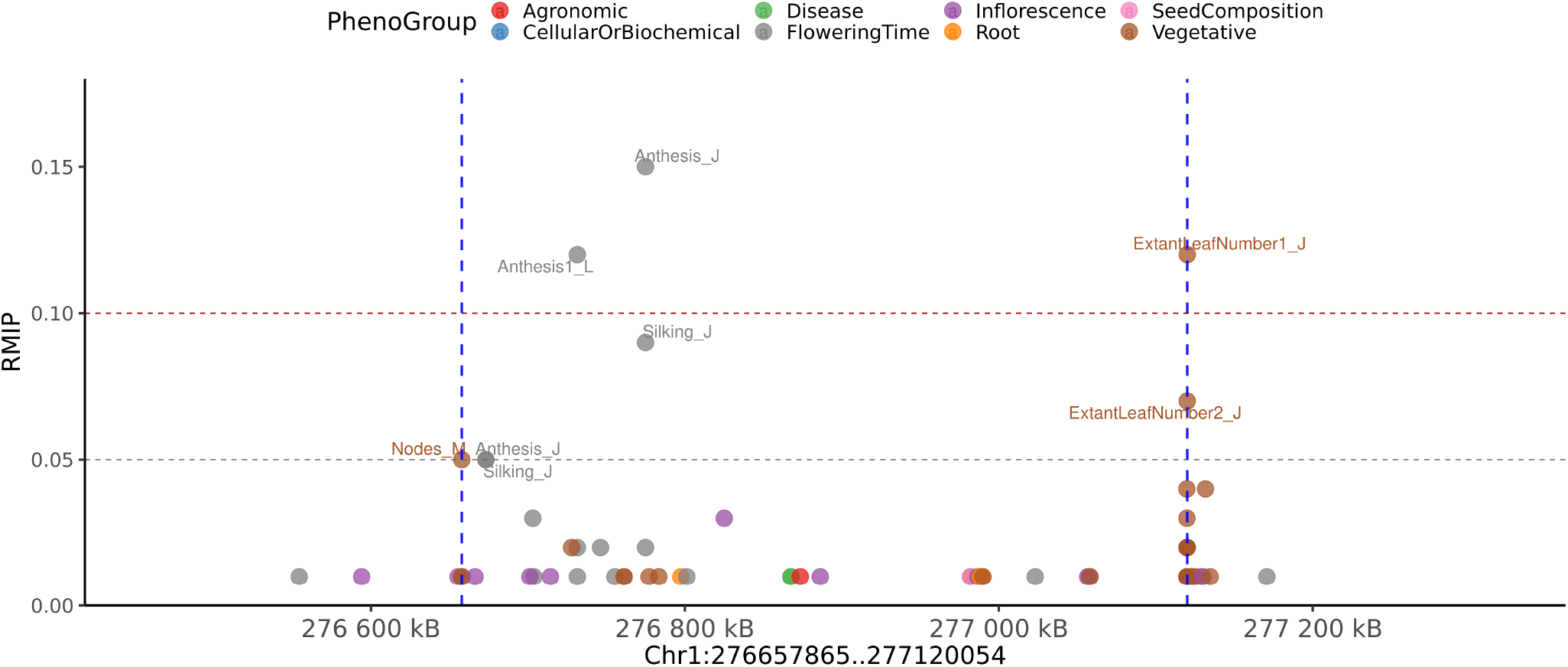
Local Manhattan plot with + /−200 kilobases of pleiotropic peak on chromosome 1 from 276,657,865 to 277,120,054. This peak is associated with the phenotypes belonging to Flowering Time and Vegetative categories. The phenotypes associated with this group are Anthesis1_L, Anthesis_J, ExtantLeafNumber1_J, ExtantLeafNumber2_J, Nodes_M, and Silking_J.

**Figure S7.**
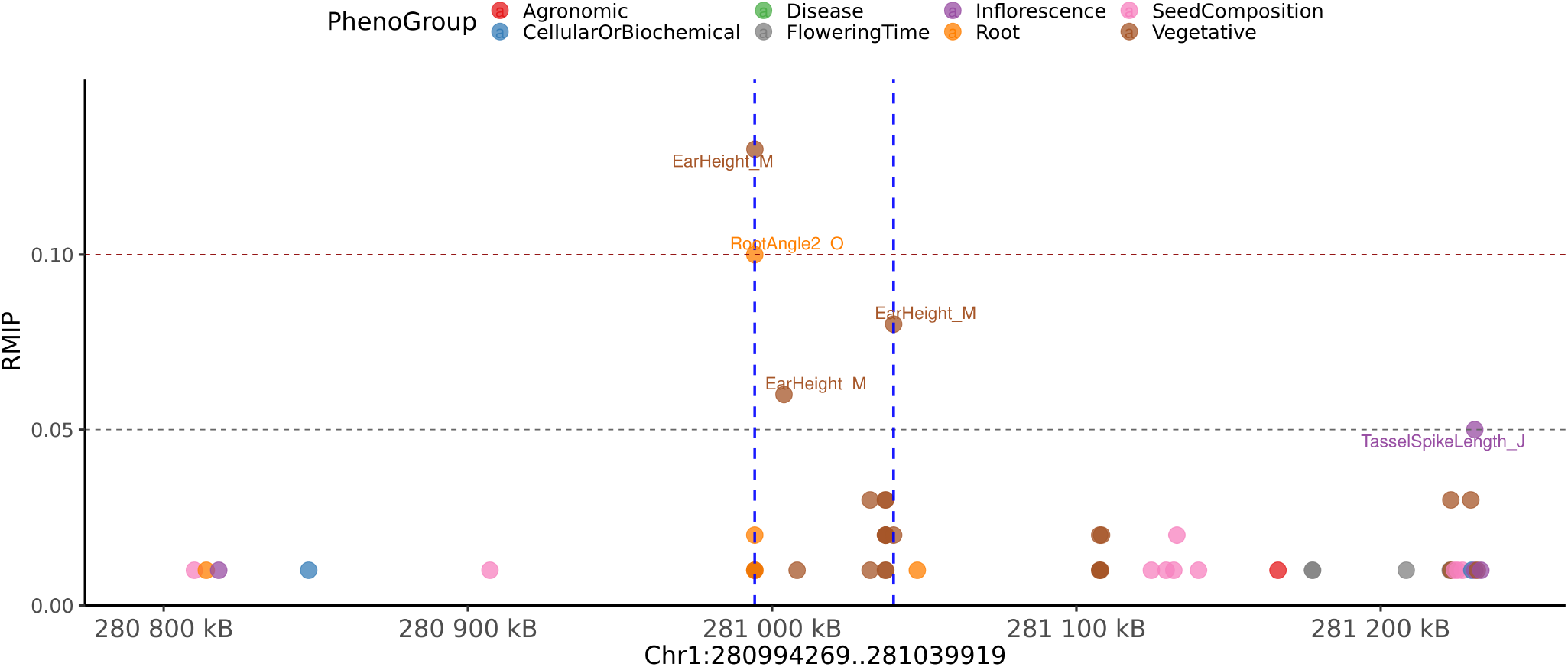
Local Manhattan plot with + /−200 kilobases of pleiotropic peak on chromosome 1 from 280,994,269 to 281,039,919. This peak is associated with the phenotypes belonging to Root and Vegetative categories. The phenotypes associated with this group are EarHeight_M and RootAngle2_O.

**Figure S8.**
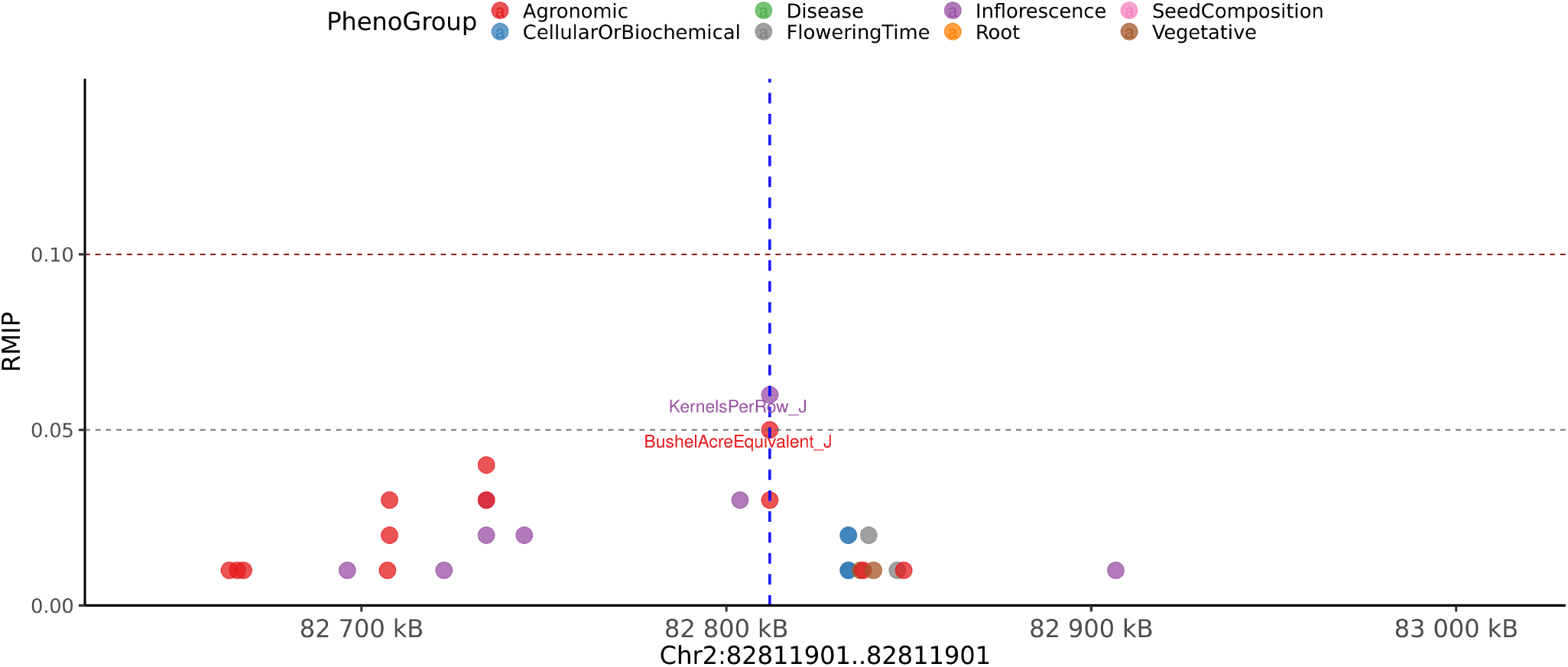
Local Manhattan plot with + /−200 kilobases of pleiotropic peak on chromosome 2 from 82,811,901 to 82,811,901. This peak is associated with the phenotypes belonging to Agronomic and Inflorescence categories. The phenotypes associated with this group are BushelAcreEquivalent_J and KernelsPerRow_J.

**Figure S9.**
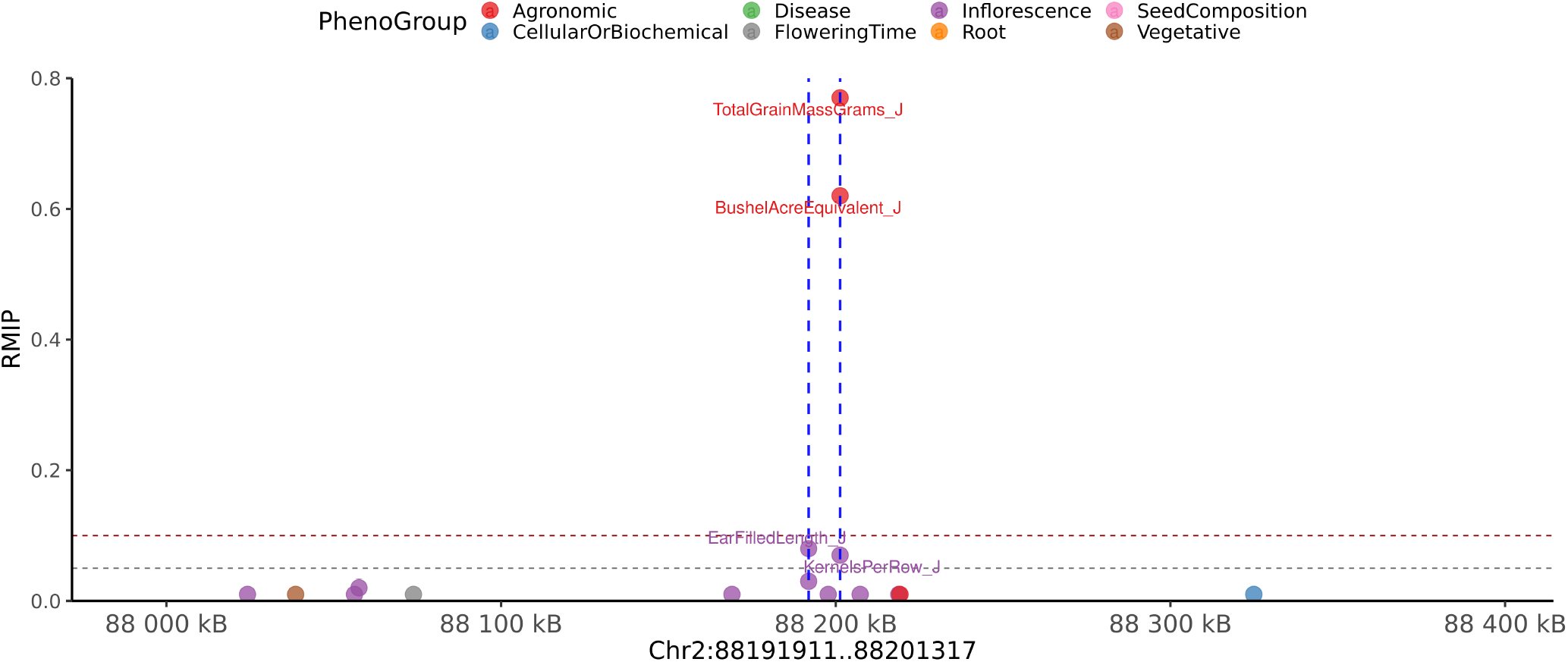
Local Manhattan plot with + /−200 kilobases of pleiotropic peak on chromosome 2 from 88,191,911 to 88,201,317. This peak is associated with the phenotypes belonging to Agronomic and Inflorescence categories. The phenotypes associated with this group are BushelAcreEquivalent_J, EarFilledLength_J, KernelsPerRow_J and TotalGrainMassGrams_J.

**Figure S10.**
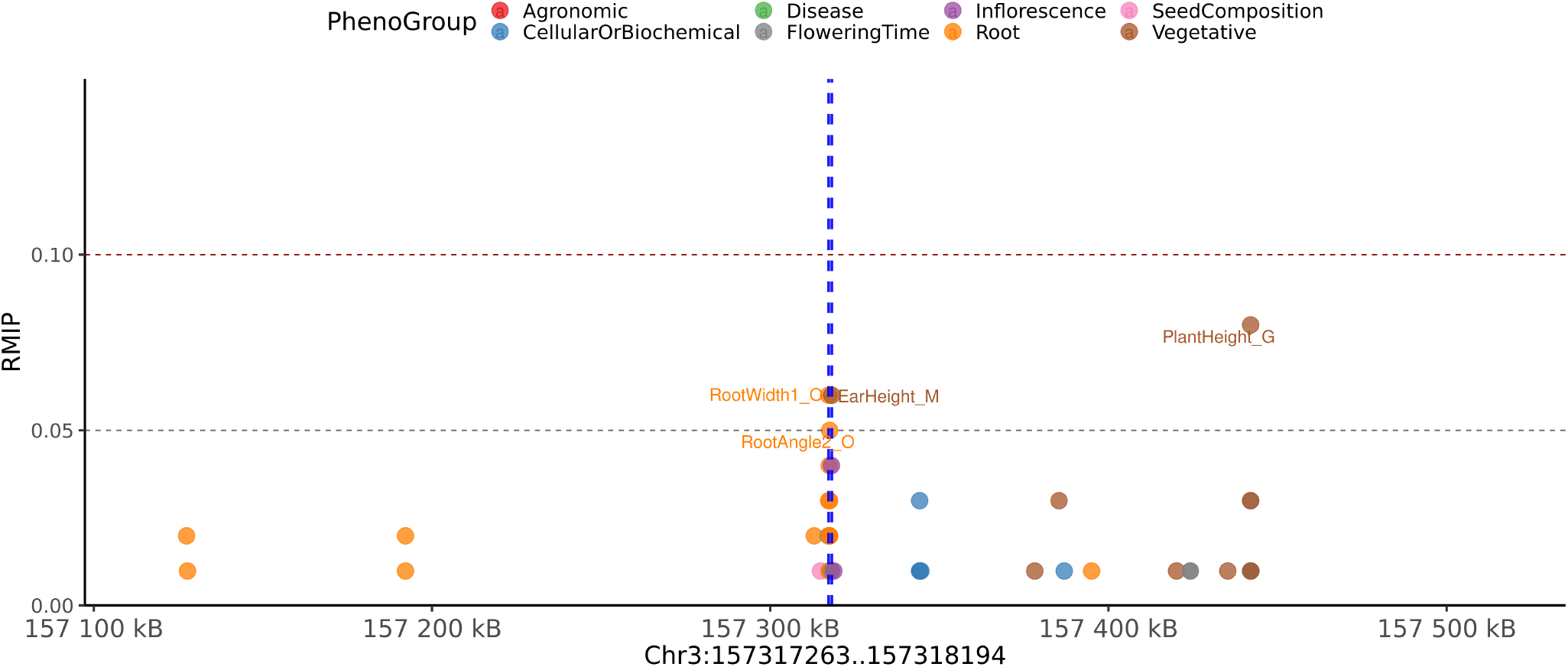
Local Manhattan plot with + /−200 kilobases of pleiotropic peak on chromosome 3 from 157,317,263 to 157,318,194. This peak is associated with the phenotypes belonging to Root and Vegetative categories. The phenotypes associated with this group are EarHeight_M, RootAngle2_O and RootWidth1_O.

**Figure S11.**
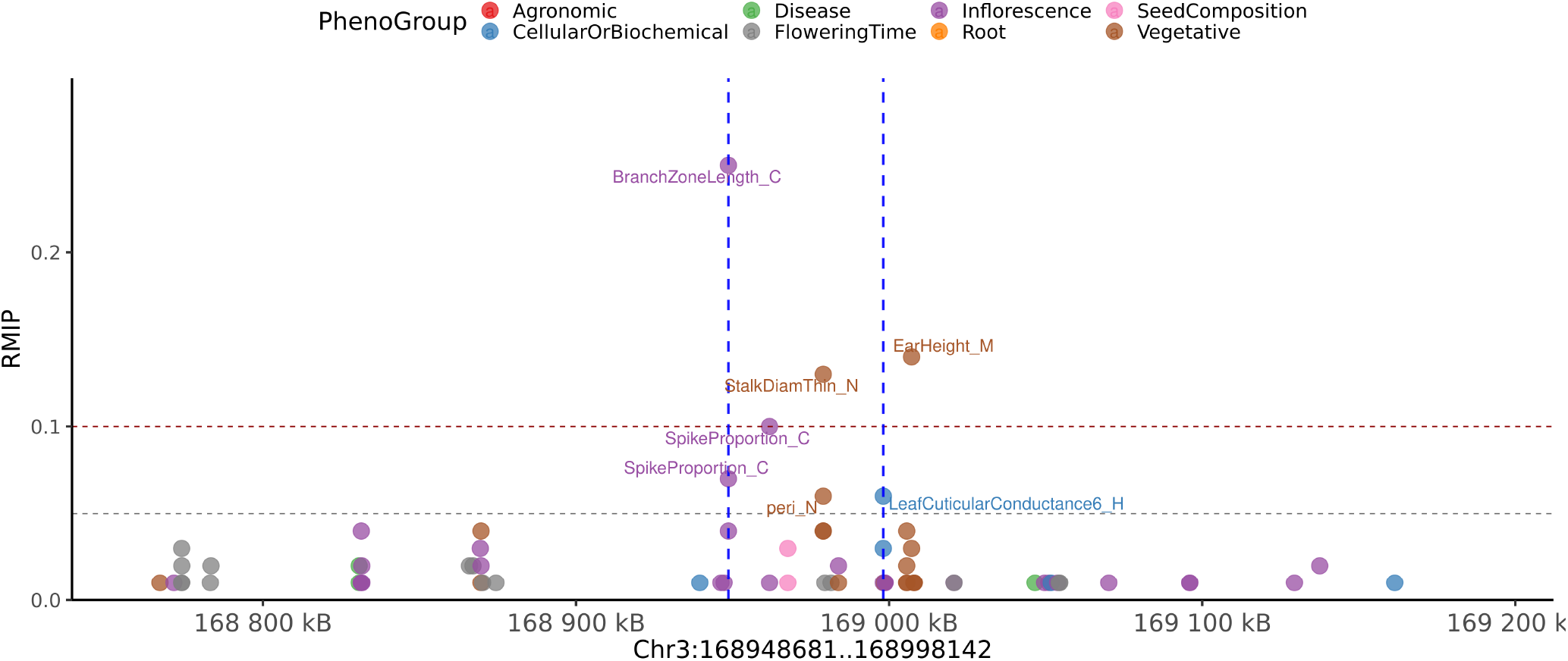
Local Manhattan plot with + /−200 kilobases of pleiotropic peak on chromosome 3 from 168,948,681 to 168,998,142. This peak is associated with the phenotypes belonging to Cellular/Biochemical, Inflorescence and Vegetative categories. The phenotypes associated with this group are BranchZoneLength_C, LeafCuticularConductance6_H, SpikeProportion_C, StalkDiamThin_N and peri_N.

**Figure S12.**
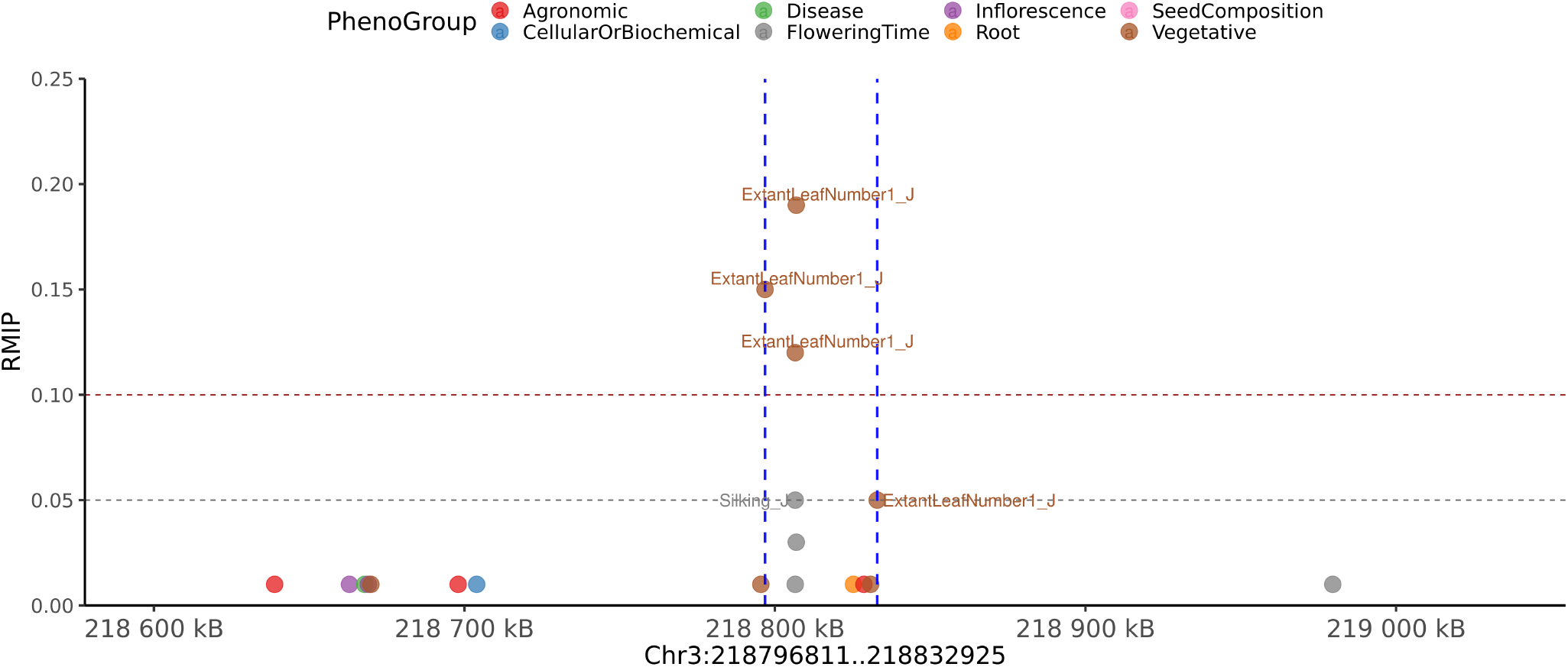
Local Manhattan plot with + /−200 kilobases of pleiotropic peak on chromosome 3 from 218,796,811 to 218,832,925. This peak is associated with the phenotypes belonging to Flowering Time and Vegetative categories. The phenotypes associated with this group are ExtantLeafNumber1_J and Silking_J.

**Figure S13.**
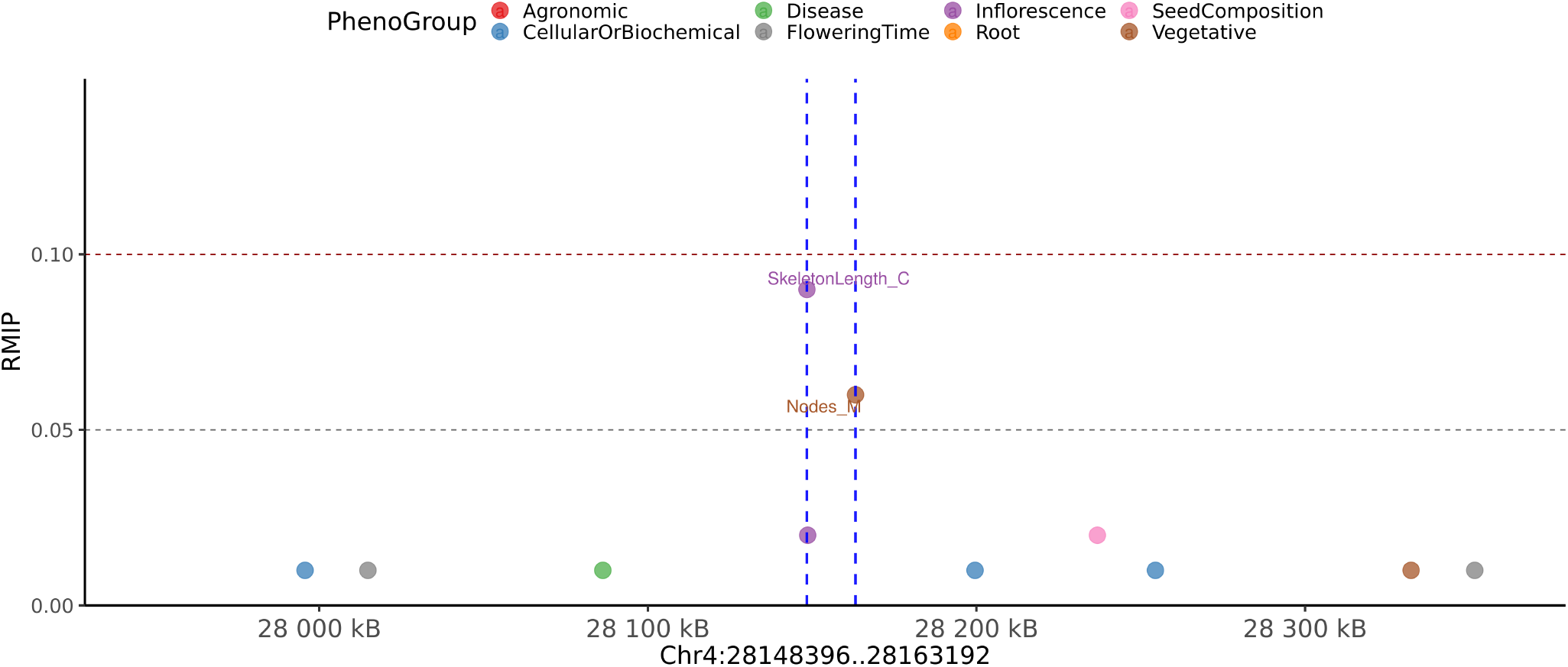
Local Manhattan plot with + /−200 kilobases of pleiotropic peak on chromosome 4 from 28,148,396 to 28,163,192. This peak is associated with the phenotypes belonging to Inflorescence and Vegetative categories. The phenotypes associated with this group are Nodes_M and SkeletonLength_C.

**Figure S14.**
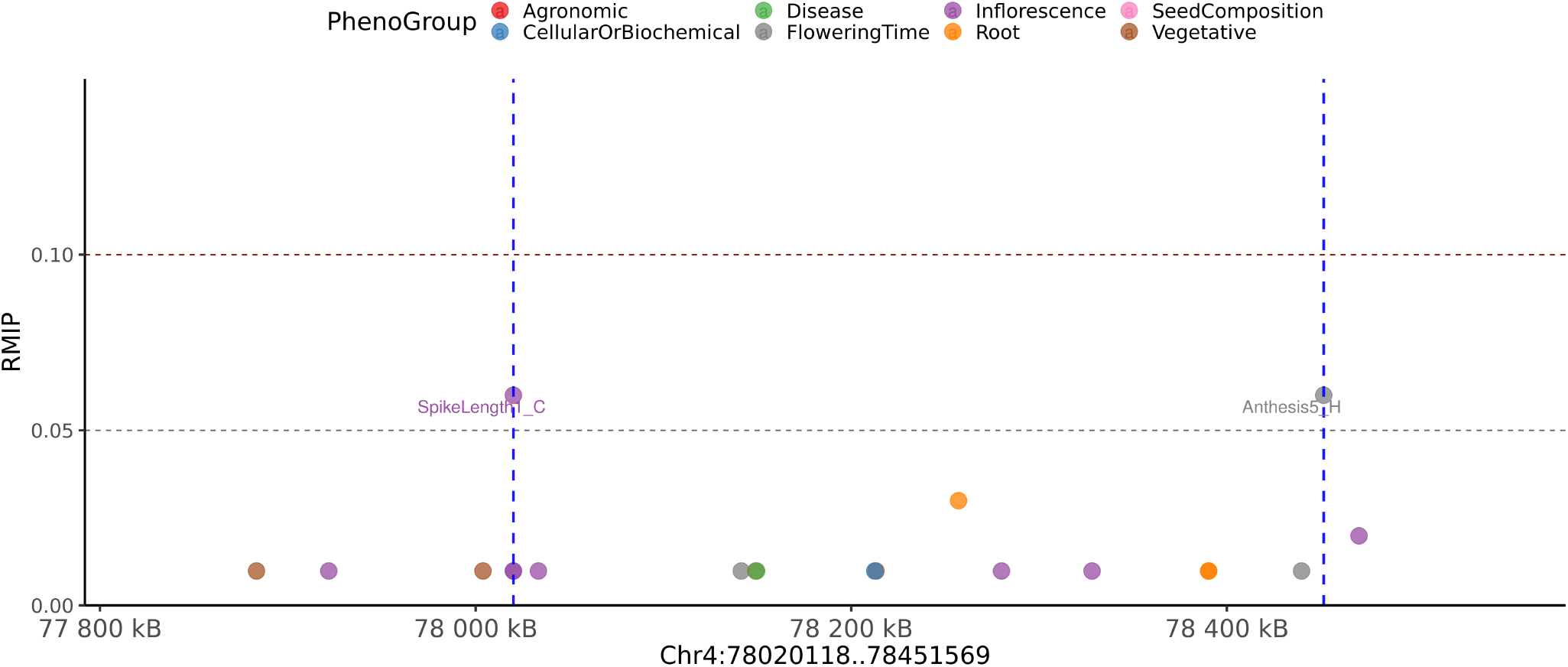
Local Manhattan plot with + /−200 kilobases of pleiotropic peak on chromosome 4 from 78,020,118 to 78,451,569. This peak is associated with the phenotypes belonging to Flowering Time and Inflorescence categories. The phenotypes associated with this group are Anthesis5_H and SpikeLength1_C.

**Figure S15.**
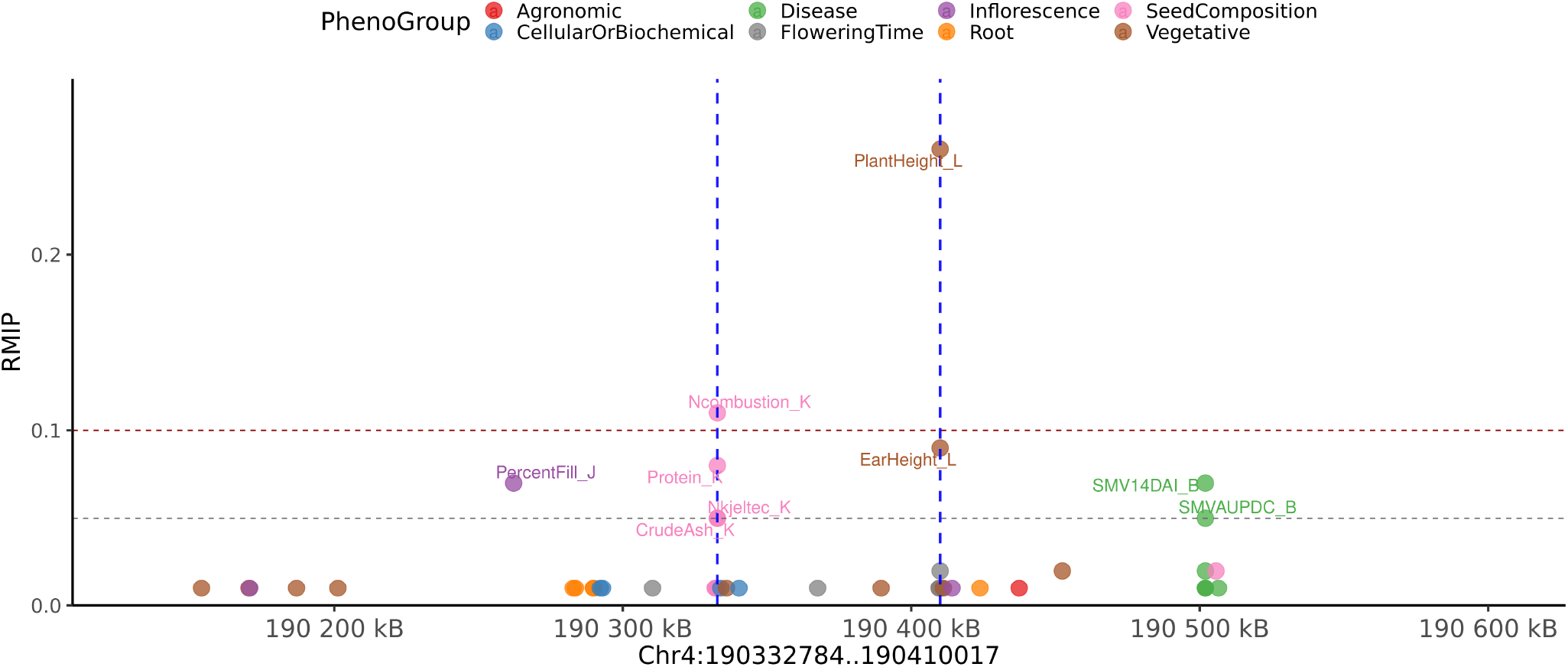
Local Manhattan plot with + /−200 kilobases of pleiotropic peak on chromosome 4 from 190,332,784 to 190,410,017. This peak is associated with the phenotypes belonging to Seed Composition and Vegetative categories. The phenotypes associated with this group are CrudeAsh_K, EarHeight_L, Ncombustion_K, Nkjeltec_K, PlantHeight_L, and Protein_K.

**Figure S16.**
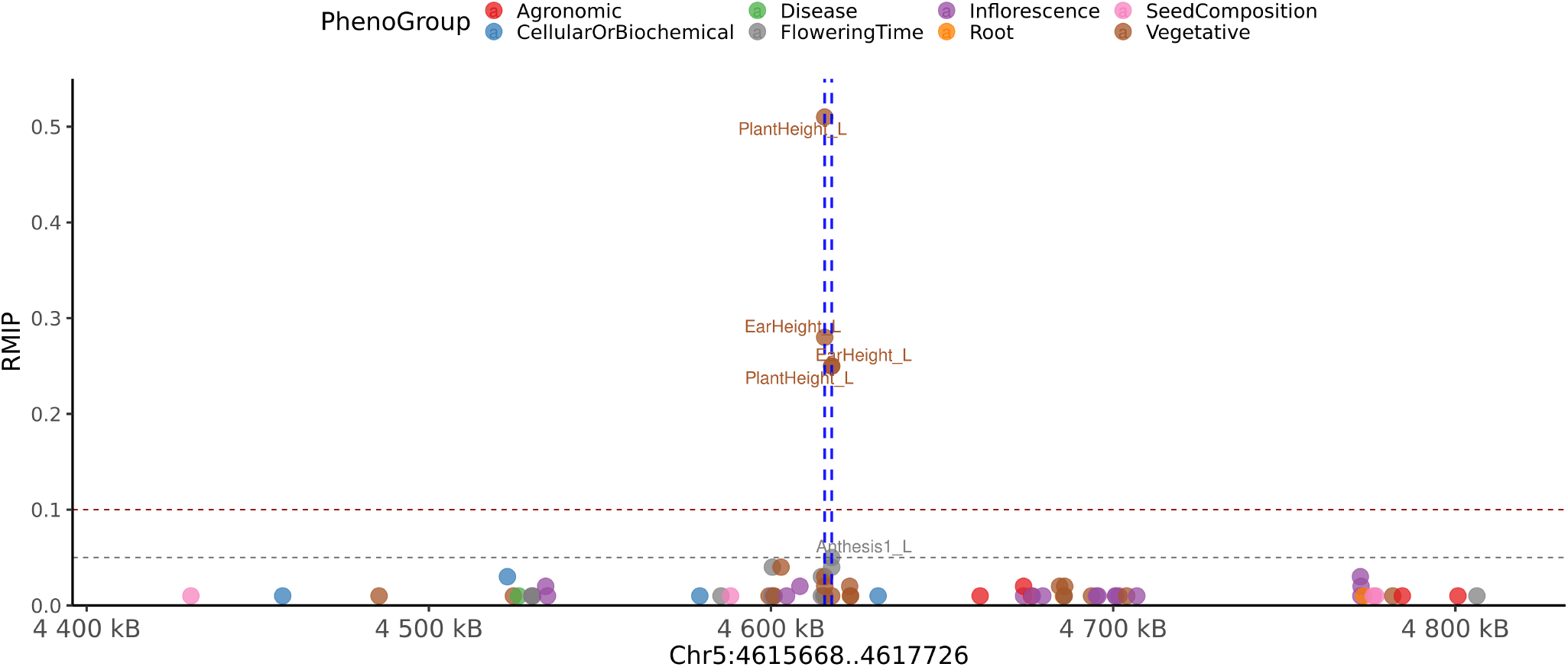
Local Manhattan plot with + /−200 kilobases of pleiotropic peak on chromosome 5 from 4,615,668 to 4,617,726. This peak is associated with the phenotypes belonging to Flowering Time and Vegetative categories. The phenotypes associated with this group are Anthesis1_L, EarHeight_L and PlantHeight_L.

**Figure S17.**
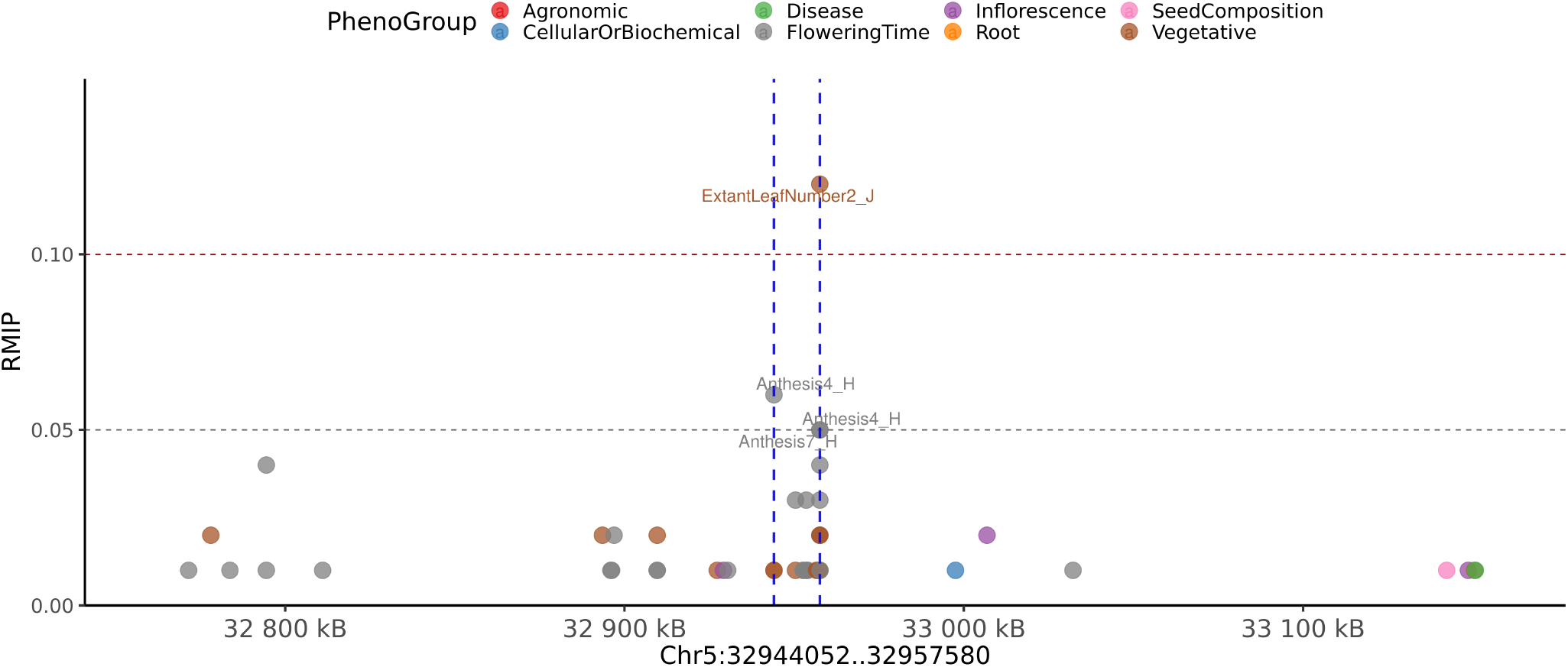
Local Manhattan plot with + /−200 kilobases of pleiotropic peak on chromosome 5 from 32,944,052 to 32,957,580. This peak is associated with the phenotypes belonging to Flowering Time and Vegetative categories. The phenotypes associated with this group are Anthesis4_H, Anthesis7_H and ExtantLeafNumber2_J.

**Figure S18.**
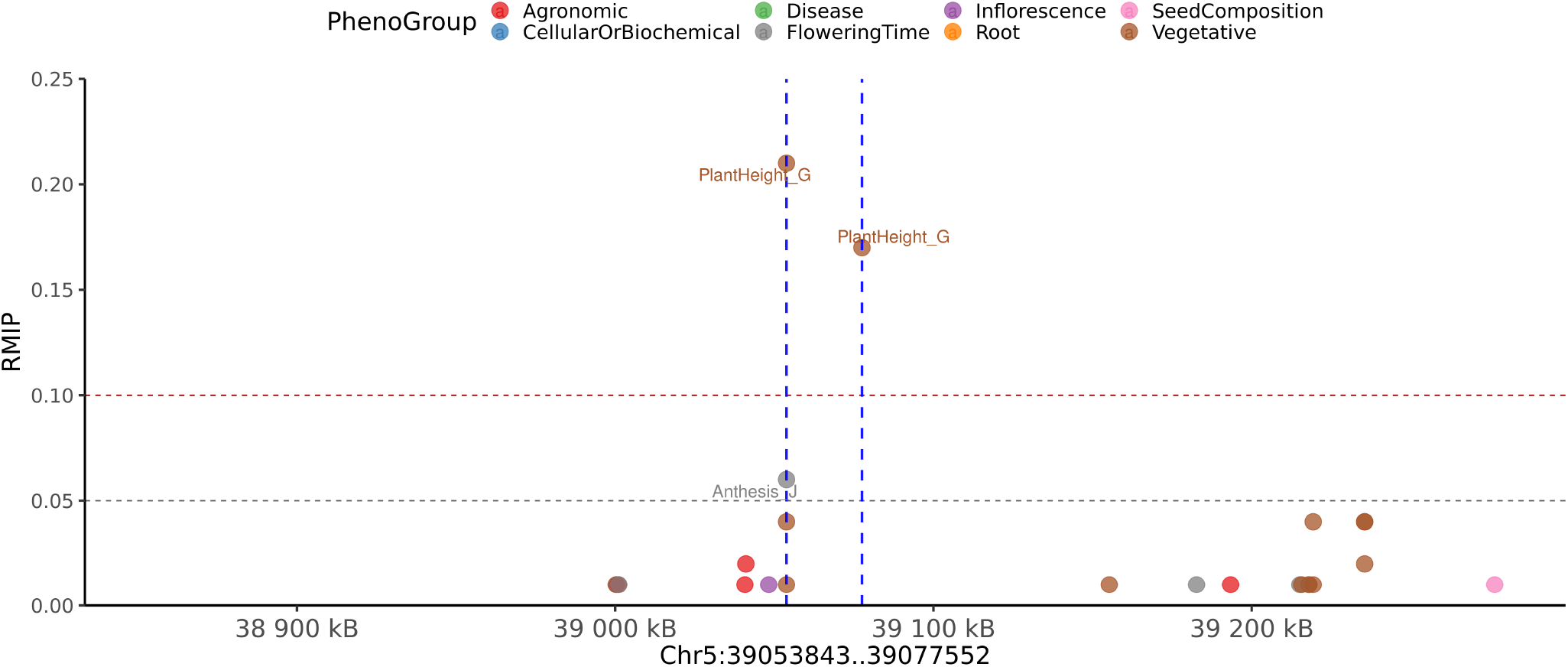
Local Manhattan plot with + /−200 kilobases of pleiotropic peak on chromosome 5 from 39,053,843 to 39,077,552. This peak is associated with the phenotypes belonging to Flowering Time and Vegetative categories. The phenotypes associated with this group are Anthesis_J and PlantHeight_G.

**Figure S19.**
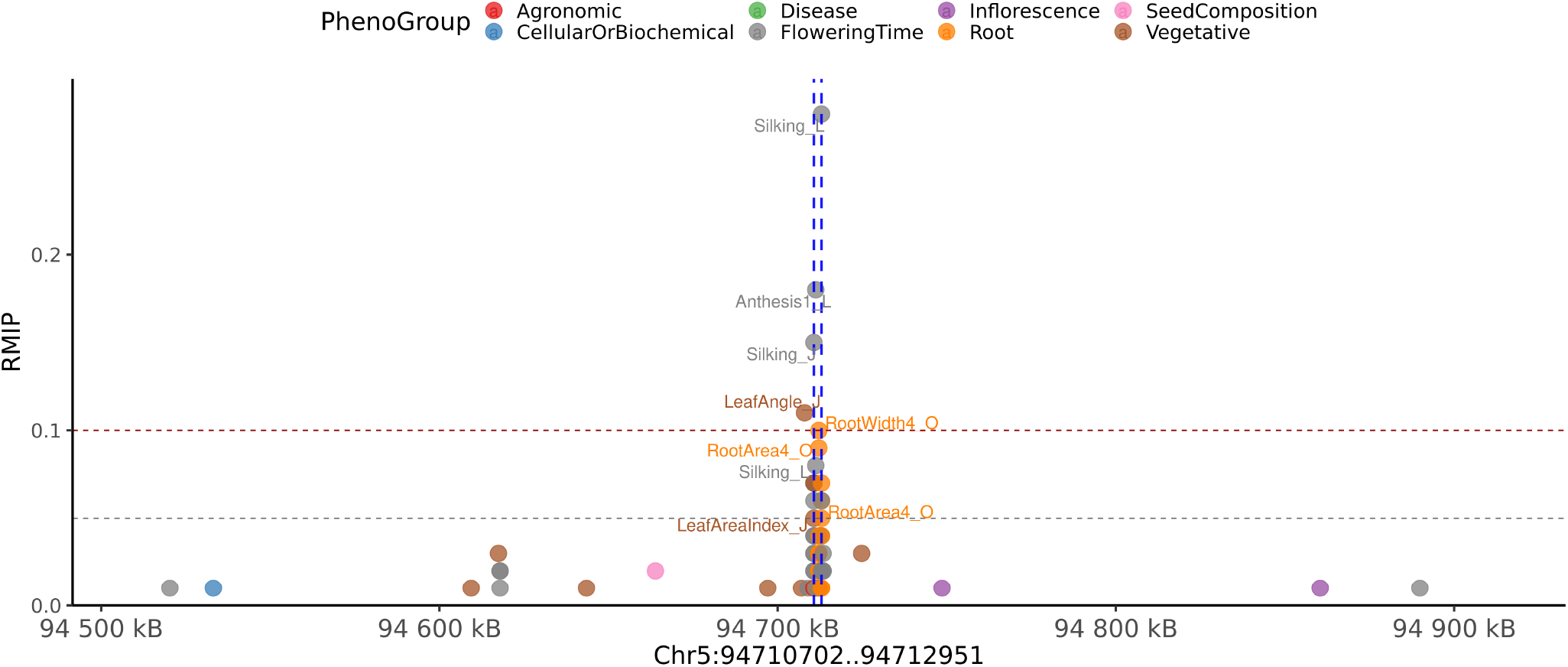
Local Manhattan plot with + /−200 kilobases of pleiotropic peak on chromosome 5 from 94,710,702 to 94,712,951. This peak is associated with the phenotypes belonging to Flowering Time, Root and Vegetative categories. The phenotypes associated with this group are Anthesis1_L, Anthesis7_H, Anthesis_A, Anthesis_J, LeafAreaIndex_J, LeafLength_J, RootArea1_O, RootArea2_O, RootArea4_O, RootWidth4_O, SilkingGDD_L, Silking_J and Silking_L.

**Figure S20.**
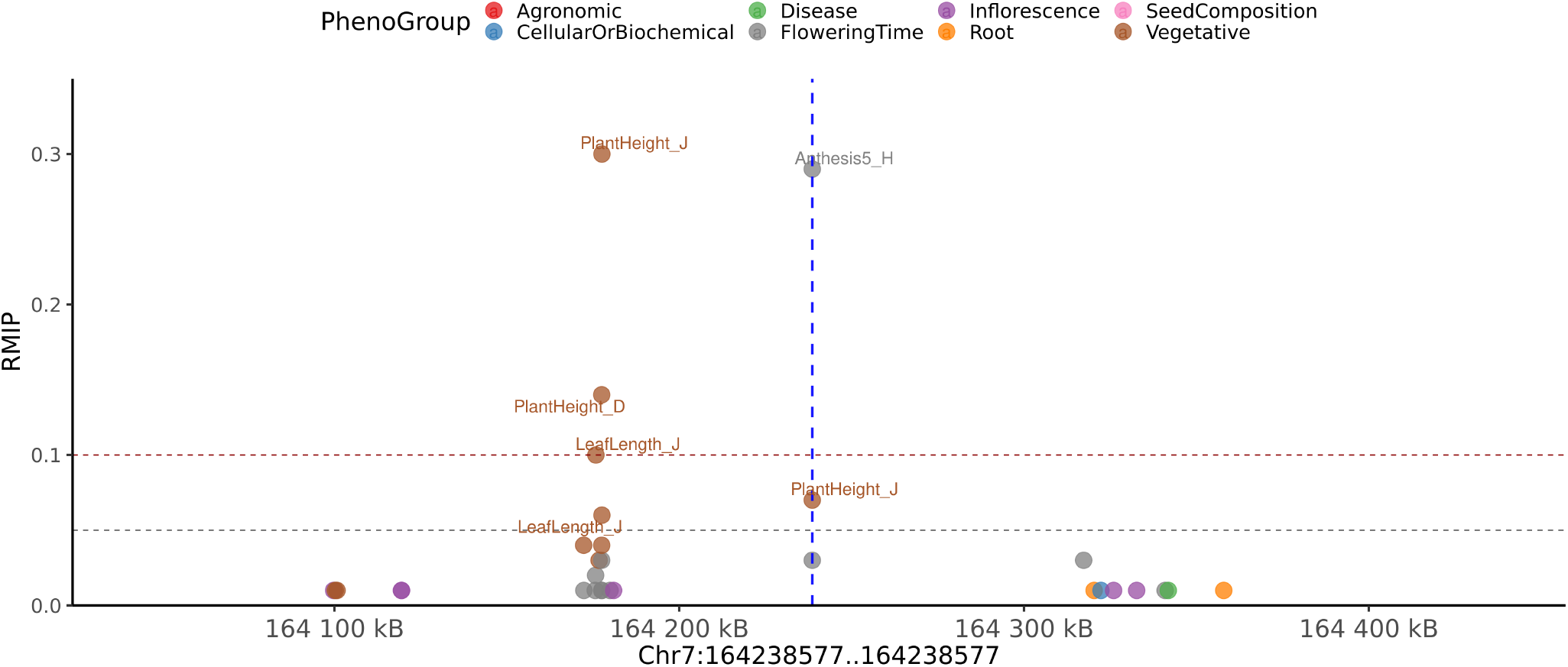
Local Manhattan plot with + /−200 kilobases of pleiotropic peak on chromosome 7 from 164,238,577 to 164,238,577. This peak is associated with the phenotypes belonging to Flowering Time and Vegetative categories. The phenotypes associated with this group are Anthesis5_H and PlantHeight_J.

**Figure S21.**
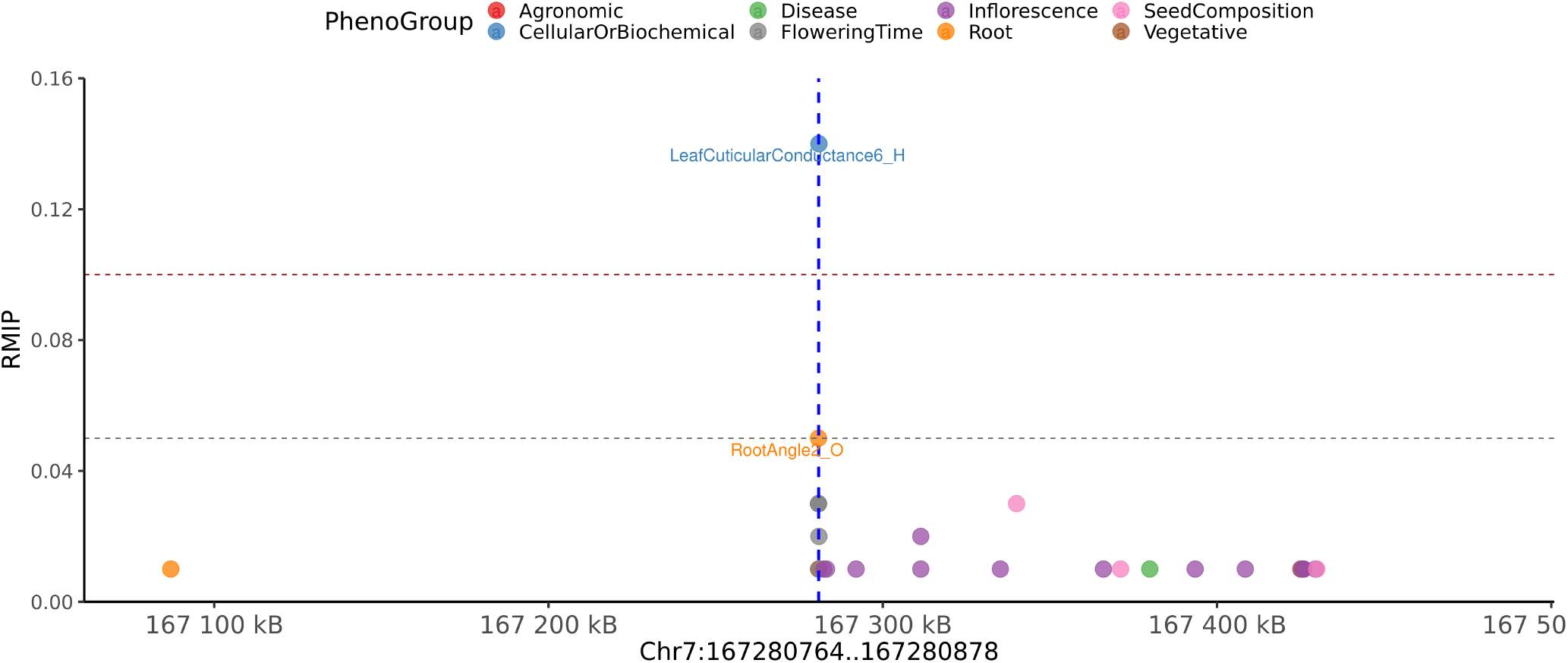
Local Manhattan plot with + /−200 kilobases of pleiotropic peak on chromosome 7 from 167,280,764 to 167,280,878. This peak is associated with the phenotypes belonging to Cellular/Biochemical and Root categories. The phenotypes associated with this group are LeafCuticularConductance6_H and RootAngle2_O.

**Figure S22.**
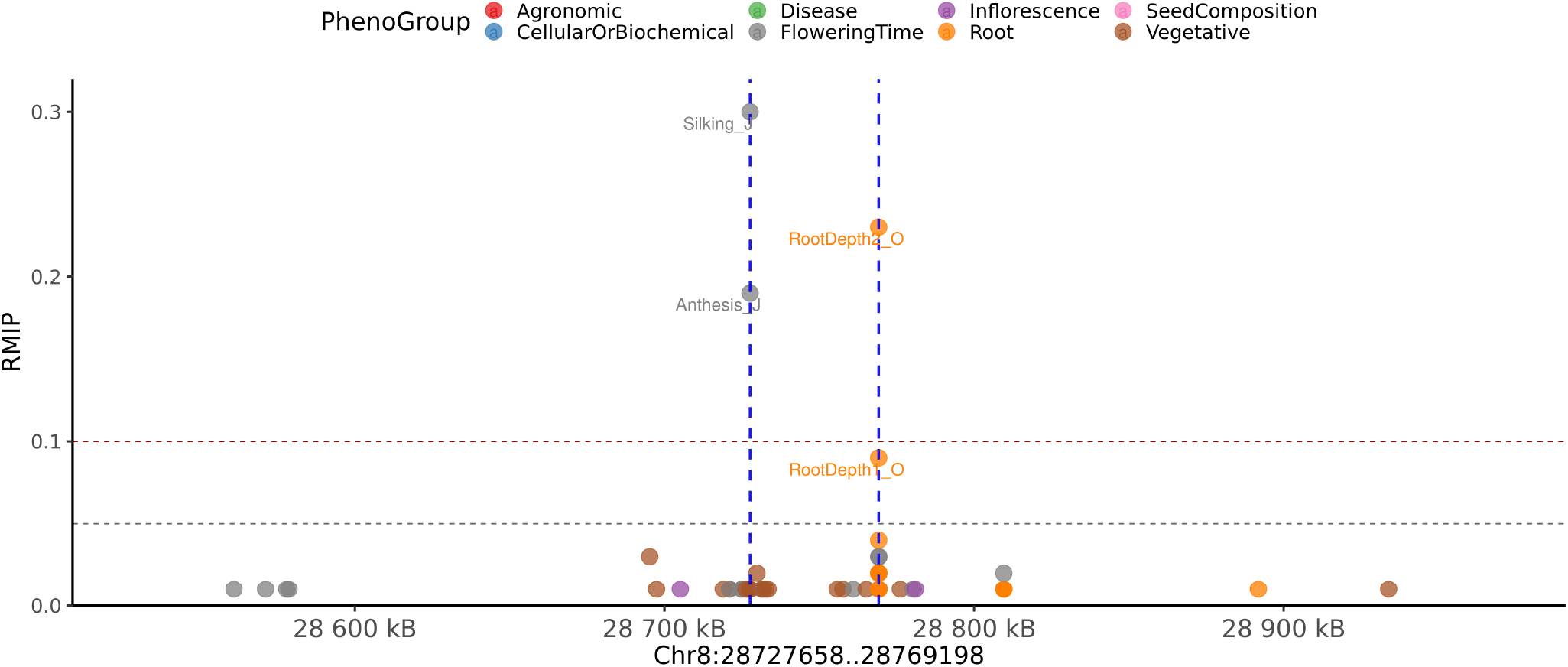
Local Manhattan plot with + /−200 kilobases of pleiotropic peak on chromosome 8 from 28,727,658 to 28,769,198. This peak is associated with the phenotypes belonging to Flowering Time and Root categories. The phenotypes associated with this group are Anthesis_J, RootDepth1_O, RootDepth2_O and Silking_J.

**Figure S23.**
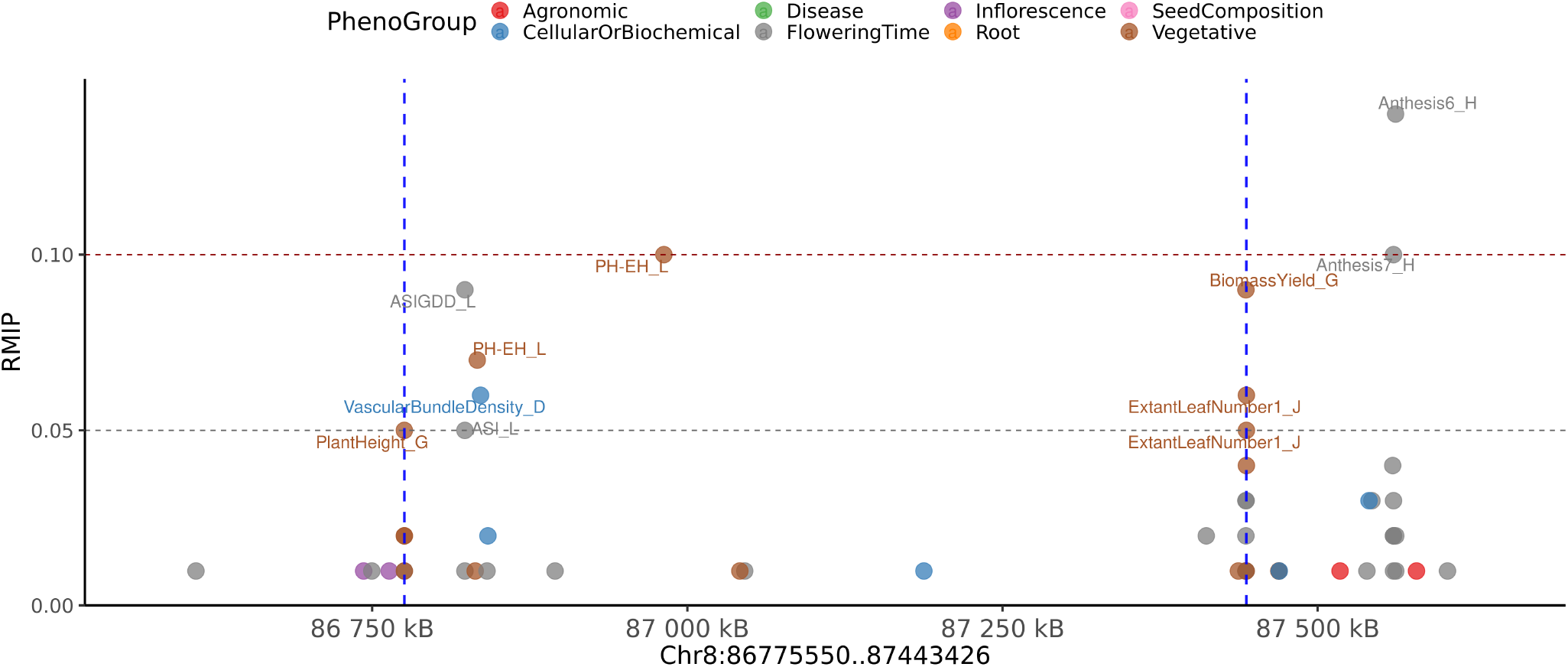
Local Manhattan plot with + /−200 kilobases of pleiotropic peak on chromosome 8 from 86,775,550 to 87,443,426. This peak is associated with the phenotypes belonging to Cellular/Biochemical and Vegetative categories. The phenotypes associated with this group are BiomassYield_G, ExtantLeafNumber1_J, PH-EH_L, PlantHeight_G and VascularBundleDensity_D.

**Figure S24.**
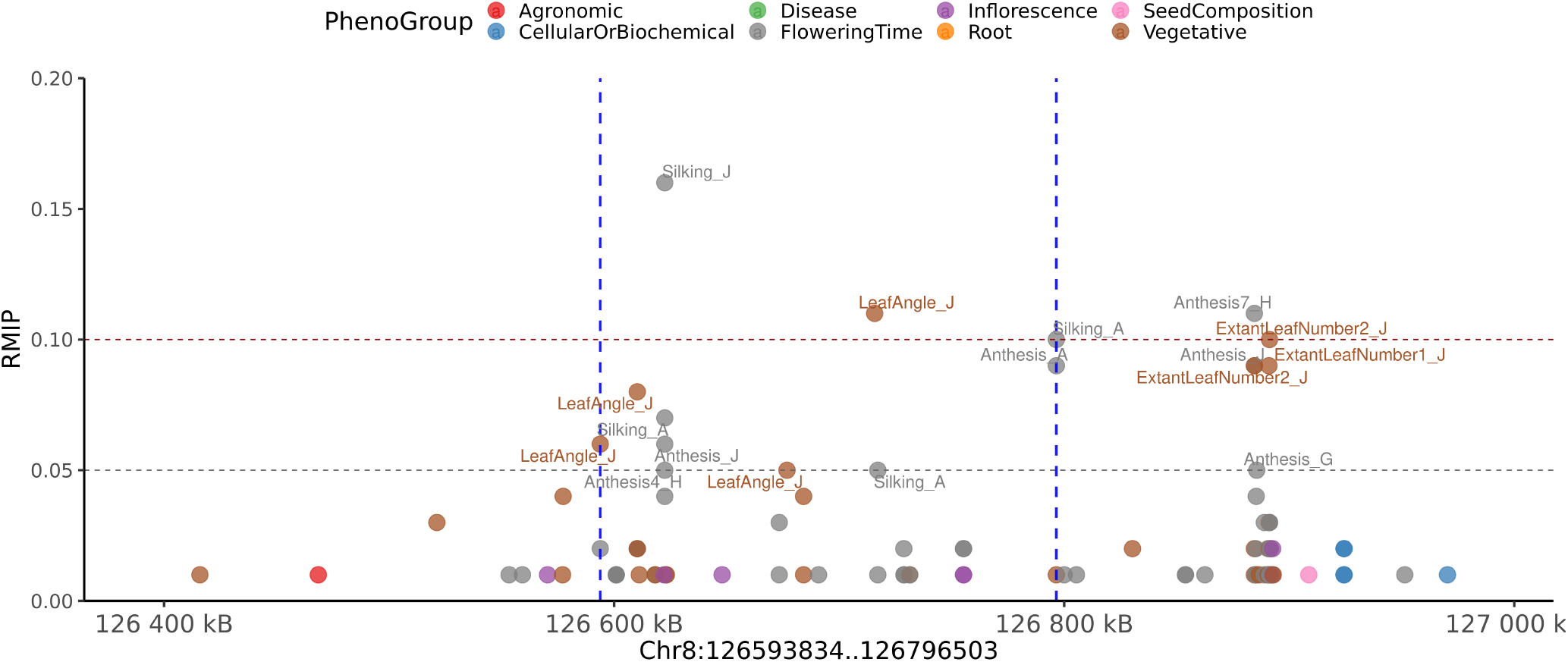
Local Manhattan plot with + /−200 kilobases of pleiotropic peak on chromosome 8 from 126,593,834 to 126,796,503. This peak is associated with the phenotypes belonging to Flowering Time and Vegetative categories. The phenotypes associated with this group are Anthesis4_H, Anthesis_A, Anthesis_J, LeafAngle_J, Silking_A and Silking_J.

**Figure S25.**
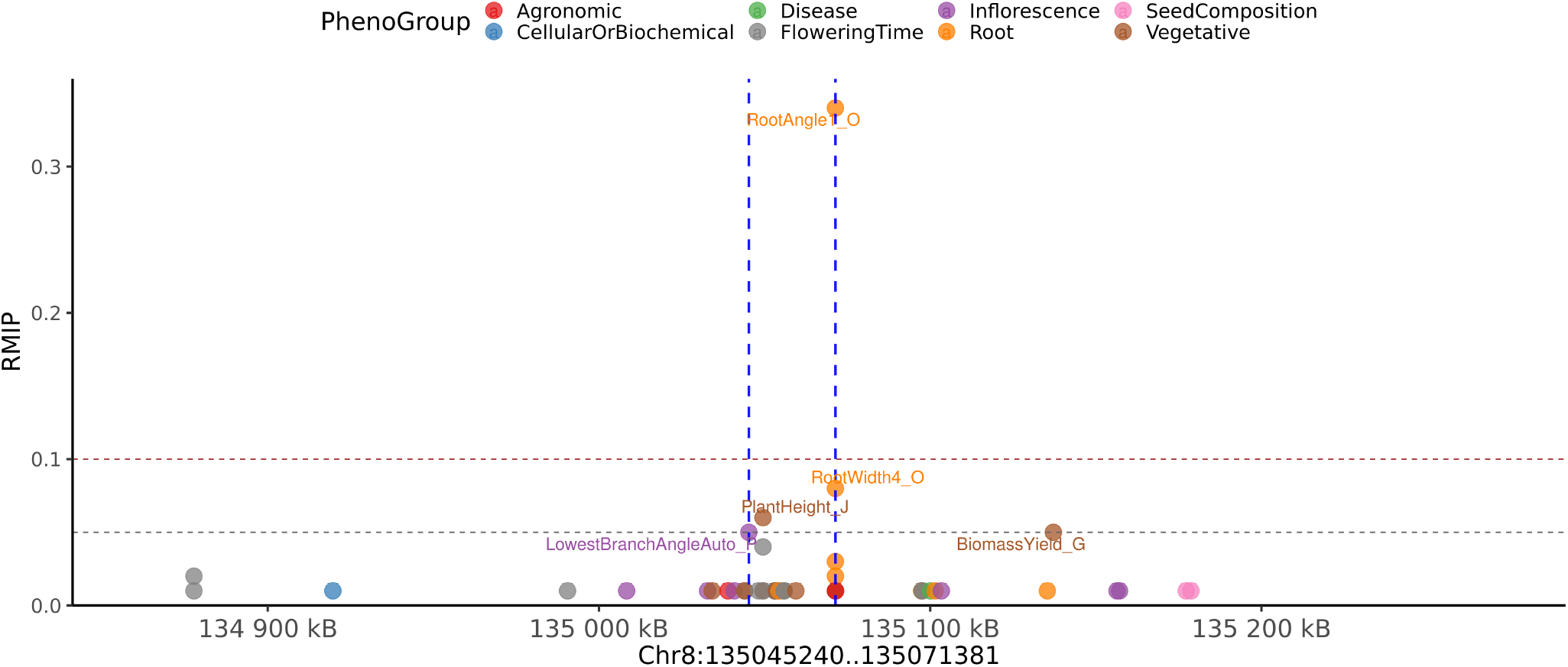
Local Manhattan plot with + /−200 kilobases of pleiotropic peak on chromosome 8 from 135,045,240 to 135,071,381. This peak is associated with the phenotypes belonging to Inflorescence, Root and Vegetative categories. The phenotypes associated with this group are LowestBranchAngleAuto_P, PlantHeight_J, RootAngle1_O and RootWidth4_O.

**Figure S26.**
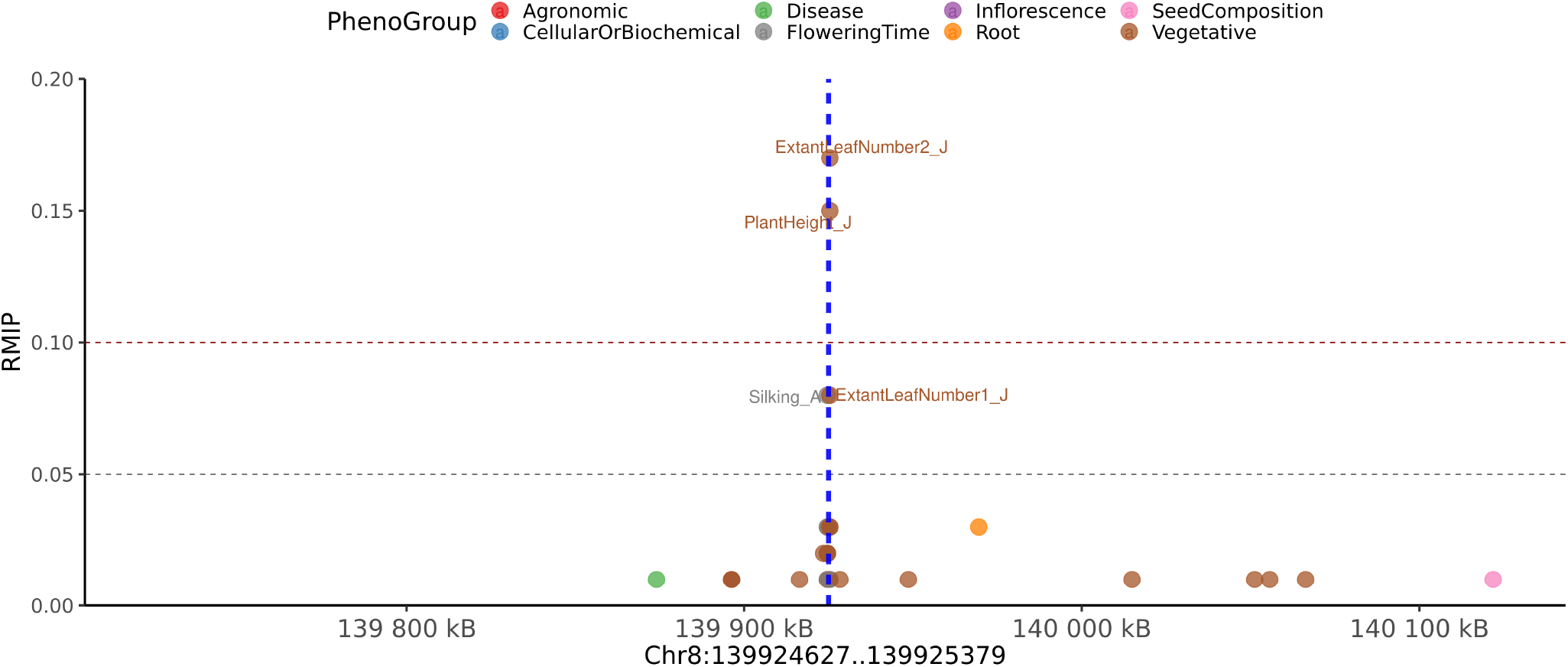
Local Manhattan plot with + /−200 kilobases of pleiotropic peak on chromosome 8 from 139,924,627 to 139,925,379. This peak is associated with the phenotypes belonging to Flowering Time and Vegetative categories. The phenotypes associated with this group are ExtantLeafNumber1_J, ExtantLeafNumber2_J, PlantHeight_J and Silking_A.

**Table S1.** Widely Used Maize Community Association Panels.

| Panel Name | Panel Abbreviation | Country | # of Accessions <sup>a</sup> | Publication |
| --- | --- | --- | --- | --- |
| Maize Association Panel | MAP | USA | 282-312 | Flint-Garcia *et al.* 2005<sup>2</sup> |
| Maize Nested Association Mapping Population | US-NAM | USA | 5000 | Jianming *et al.* 2008<sup>65</sup> |
| Ames Inbred Lines | NCRPIS | USA | 2815 | Romay *et al.* 2013<sup>33</sup> |
| Maize Shoot Apical Meristem | SAM | USA | 369 | Leiboff *et al.* 2015<sup>3</sup> |
| Wisconsin Diversity Panel | WiDiv | USA | 503-942 | Hanesy *et al.* 2011<sup>4</sup>, and Mazaheri *et al.* 2019<sup>6</sup> |
| China Nested Association Mapping Population | CN-NAM | China | 1971 | Li *et al.* 2013<sup>66</sup> |
| Global Diverse Lines |  |  | 527 | Yang *et al.* 2011<sup>67</sup> |
| Dent and Flint Panel |  | Europe | 306 + 292 | Revilla *et al.* 2014<sup>68</sup> |
| European Flint Collection |  | Europe | 1191 | Gouesnard *et al.* 2017<sup>69</sup> |
| European Flint Nested Association Mapping | EUNAM-Flint | Europe | 811 | Lehermeier *et al.* 2014<sup>70</sup> |
| European Dent Nested Association Mapping | EUNAM-Dent | Europe | 841 | Lehermeier *et al.* 2014<sup>70</sup> |

<sup>a</sup> The number of lines have changed over time due to the loss of some genotypes and/or the addition of others.

**Table S2. Trait values for all phenotypes per accession and associated information per accession.** Provided as included excel file.

**Table S3. Phenotypes and associated information.** Provided as included excel file.

**Table S4. The locations of GWAS peaks, the traits associated with each peak and the genes adjacent to each peak.** Provided as included excel file.

**Table S5. Individual GWAS hits.** Provided as included excel file.

